# Harmful algal bloom species *Microcystis aeruginosa* releases thiamin antivitamins to suppress competitors

**DOI:** 10.1101/2025.04.17.649140

**Authors:** Mohammad Yazdani, Christopher P. Suffridge, Fangchen Liu, Cait M. Costello, Zhiyao Zhou, Gillian St. John, Ruchika Bhawal, Sheng Zhang, Geoffrey W. Coates, Mingming Wu, Beth A. Ahner

## Abstract

In environmental ecosystems, vitamin concentrations are often exceedingly low (1, 2) and auxotrophy, or reliance on exogenous vitamin or vitamin precursors, is widespread (3–5). We show here that the widespread harmful algal bloom (HAB) species *Microcystis aeruginosa,* threatening freshwater aquatic ecosystems globally, releases a complex mixture of thiamin antivitamins, including bacimethrin and methoxythiamin, which induce thiamin deficiency in the benign model green alga *Chlamydomonas reinhardtii*. Putative biosynthetic genes for bacimethrin were upregulated in *M. aeruginosa* when grown in co-culture resulting in greater production of bacimethrin. Bacimethrin, methoxythiamin, oxidized forms of thiamin and methoxythiamin, and a novel structural homolog of bacimethrin were all found at elevated levels in the co-culture exometabolome extracts and were all inhibitory to the growth of *C. reinhardtii* individually at very low concentrations and as a mixture in culture medium extracts. The thiamin-requiring mutant *C. reinhardtii*, CC-25, was much more sensitive to bacimethrin and methoxythiamin than the wildtype. Thiamin addition largely rescued the inhibitory effects of exposure to antivitamins in both the wildtype and mutant strain. Finally, we determined that bacimethrin is present in aquatic environments and is elevated during *Microcystis* blooms. Thus, allelopathic suppression of competitors, particularly those that are auxotrophic for thiamin, by *M. aeruginosa* via the production of antivitamins in environments where thiamin availability is low, could help this species to become dominant and form blooms.

---

Freshwater and marine ecosystems worldwide are increasingly threatened by harmful algae blooms (HABs) which can cause widespread fish, bird, and mammal death as well as major disruptions to drinking water supply and recreation (6, 7). HABs occur when toxin-producing species of cyanobacteria or microalgae out-compete non-toxic species for macronutrients, light, and other resources. Eutrophication is cited as a leading cause of HABs and climate change appears to be increasing the scope and frequency of blooms, however our understanding of factors leading to bloom formation and their persistence is far from complete (7–9).

Microbes in communities from diverse environments, including aquatic ecosystems where HABs occur, are known to exchange important cellular metabolites (10), including vitamins (11). Indeed, exchange of vitamins is so prevalent that auxotrophy, or the complete reliance on an exogenous source of vitamin or vitamin precursors, is common in evolutionarily diverse species (3–5). In aquatic ecosystems, dissolved concentrations of required vitamins, such as thiamin, are also exceedingly low (1, 2) and the bioavailability of vitamins has been hypothesized to influence plankton species diversity and succession in seawater (1, 12, 13).

Also prevalent in microbial communities are allelopathic interactions, where one organism suppresses another by releasing an inhibitory chemical compound (14). Cyanobacterial species, such as *Microcystis aeruginosa* which is responsible for toxic blooms in many freshwater systems, are known to release cyanotoxins (15) or lipophilic metabolites (16, 17) to inhibit benign green algae that naturally compete for light and nutrients. However, evidence from field studies confirming that allelopathic interactions are instrumental in species succession or bloom formation is limited (17) and a role for allelopathy in dilute and turbulent marine ecosystems has been disputed (18). Due to the limited availability of vitamins in surface water and the known presence of high affinity vitamin transport proteins(19), allelopathic chemicals with structural similarity to vitamins or their precursors may be very effective inhibitors of microbial activity.

Chemicals that are inhibitory or toxic because they mimic vitamins are called antivitamins. Most research on antivitamins has been focused on drug discovery (20) from which it is known that multiple bacterial species synthesize thiamin antivitamins (21–23). Given that the highly successful freshwater cyanobacteria *M. aeruginosa* can synthesize its own thiamin, we hypothesize that an overlooked secret to its success might involve the production and release of thiamin antivitamins to which it is more resistant than its competitors (Fig 1A). Here we used a co-culture, including *M. aeruginosa* and the benign model green algae *Chlamydomonas reinhardtii,* to show that thiamin antivitamins are produced by *M. aeruginosa,* and that the presence of the competing algae influences antivitamin accumulation. We also document elevated levels of one antivitamin during a *Microcystis* bloom and, using a *C. reinhardtii* mutant, confirm that species unable to make their own thiamin are likely to be particularly sensitive to this type of allelopathy.

**Fig. 1.**
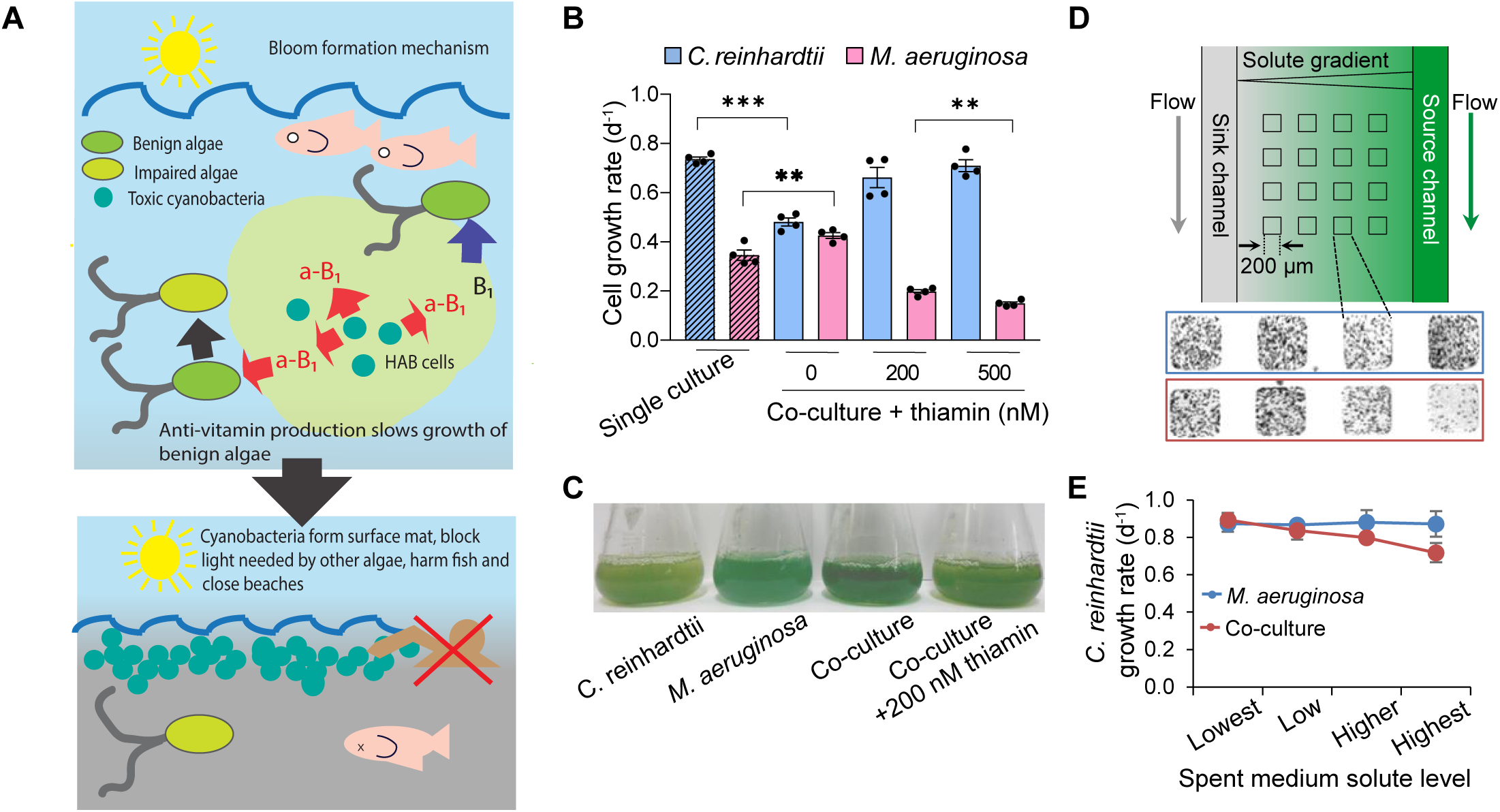
(A) Illustration of proposed thiamin antivitamin (a-B1) allelopathic mechanism leading to HABs. Dissolved vitamin B1 is very low but is required by algae; a-B1 is nearly identical in structure but toxic. HAB cells release a-B1 and other chemicals to slow the growth of benign algae. Over time HAB forms surface mat, shades other algae, produces toxins closing beaches and harms fish populations. (B) Growth rate of *C. reinhardtii* and *M. aeruginosa* in single culture, co-culture, and co-culture with thiamin. (C) At t = 10 d, from left to right, single cultures of *C. reinhardtii* and *M. aeruginosa*, and the co-cultures without (blue-green) and with thiamin (yellow-green). (D) Schematic of the microfluidic device used to generate gradient of exudates from spent medium in replicate microhabitats. Epi-fluorescent microscope images of cells in microhabitats at t =3 d from across solute gradients. (E) *C. reinhardtii* growth rate in individual microhabitats. For B and E data are means ± SEM, with n ≥ 3. Asterisks indicate significance in Student’s t-test (\*\**P* < 0.01, \*\*\**P* < 0.001).

## Results

### Thiamin Rescues Growth Inhibition of *C. reinhardtii* by *M. aeruginosa*

Cyanobacteria produce many antimetabolites, some of which are toxic to photosynthetic eukaryotes (15–17, 24). Accordingly, when we cultured *M. aeruginosa* together with *C. reinhardtii* in minimal medium BG-11, it was not surprising that the latter grew 35% slower than when grown alone (Fig. 1*B*). In contrast, *M. aeruginosa* grew at a rate that was 23% higher than when grown alone (Fig. 1*B*) and ultimately dominated the co-culture after 10 days (Fig. 1*C*). To determine whether exudates were responsible for the observed inhibition, we evaluated the growth of *C. reinhardtii* in a microfluidic device that can be used to expose cells to a steady state gradient of solutes (25) (Fig. 1*D*). *C. reinhardtii* cells were seeded in an array of microhabitats patterned into an agarose membrane and a chemical gradient was established via diffusion by pumping spent and fresh medium into the source and sink channels respectively. After three days, microscopy showed significantly fewer *C. reinhardtii* cells in the microhabitats exposed to higher solute levels from the co-culture spent medium (rightmost microhabitat in red box, Fig. 1*D*) than those closer to the sink channel. The growth rate was also the lowest in these microhabitats (Fig. 1*E*). In contrast, the growth rate of *C. reinhardtii* was the same in all the microhabitats exposed to a gradient of solutes from the spent medium of *M. aeruginosa* culture (Fig. 1*E*).

Given that one previously characterized allelopathic interaction induced oxidative stress (17), we added thiamin, a vitamin known to enhance plant tolerance to oxidative stress (26), to a subsequent set of co-cultures. In the presence of thiamin, the growth rate of *C. reinhardtii* fully recovered and, in contrast, that of *M. aeruginosa* decreased (Fig. 1*B*), resulting ultimately in a co-culture dominated by *C. reinhardtii* instead of *M. aeruginosa* to (Fig. 1*C*). Thiamin addition to single cultures of *M. aeruginosa* did not cause inhibition whereas *C. reinhardtii* growth was slightly enhanced (Fig. S1). Given the strong link we observed between thiamin addition and co-culture domination, we decided to test an alternate hypothesis regarding thiamin’s role in mitigating allelopathy: the production of antivitamins by *M. aeruginosa*.

### *M. aeruginosa* Produces Thiamin Antivitamins

Spent medium from the co-culture and from single cultures of both species was lyophilized and the chemical residuals were resuspended in methanol to further study the composition of the *M. aeruginosa* exudates. The co-culture methanol extract, like the spent medium in the microfluidic device, suppressed *C. reinhardtii* growth by up to 50% of controls, whereas *M. aeruginosa* single culture extract only reduced its rate by 20% (Fig. 2*A*); the co-culture extract did not affect *M. aeruginosa* growth (Fig. S2). When thiamin was added with the co-culture extract, significantly less inhibition of *C. reinhardtii* was observed (Fig. 2*A*). Non-targeted liquid chromatography mass spectrometry (LC-MS/MS) analysis of methanol extracts was used to characterize the set of metabolites shed by cells into the growth medium, i.e., the exometabolomes (Dataset S1). An orthogonal projection to latent structures discriminant analysis (OPLS-DA) plot of the 1145 annotated compounds in the exometabolomes revealed that the co-culture exometabolome overlaps with the two derived from the individual cultures but is not simply a composite of the two (Fig. 2*B*).

**Fig. 2.**
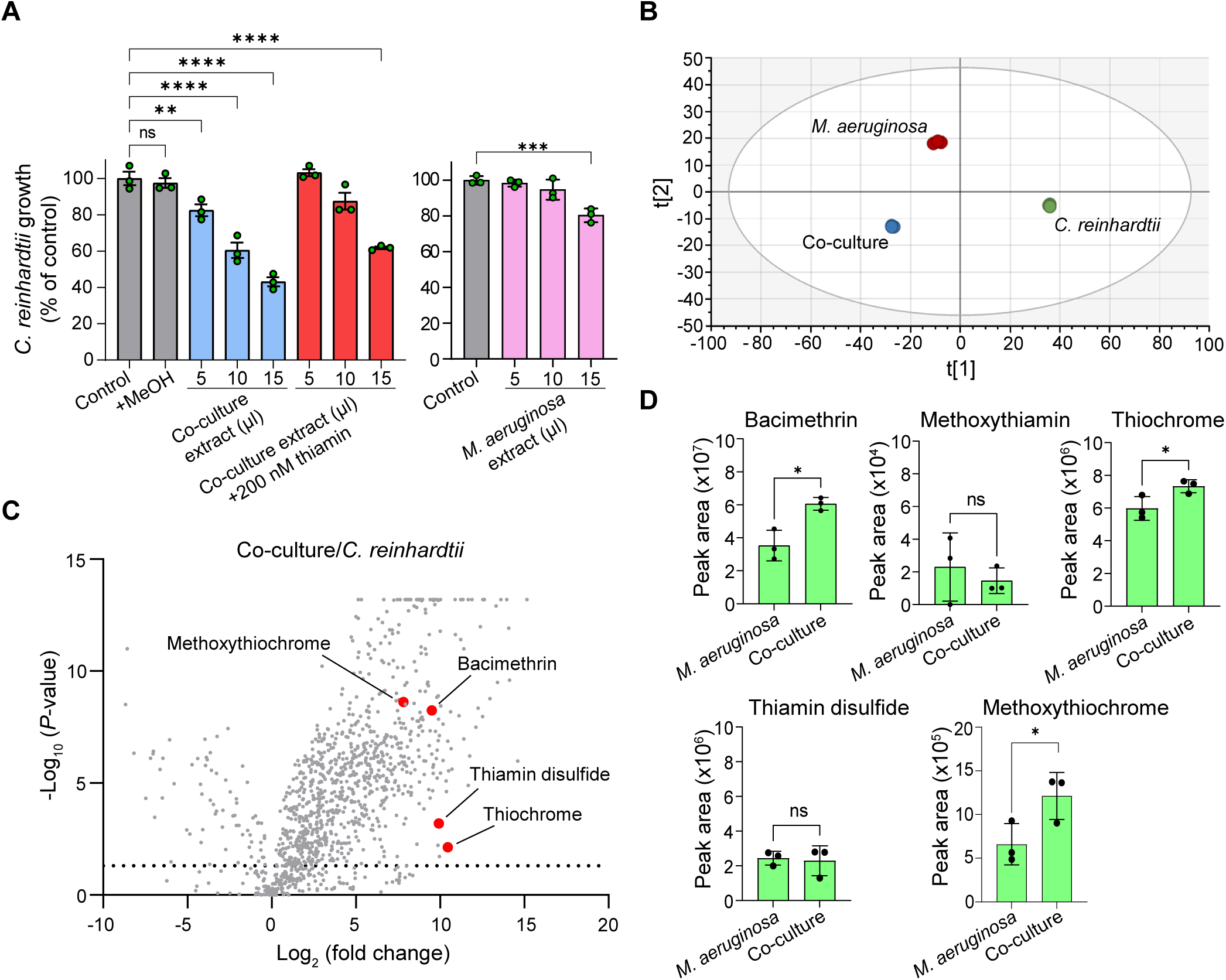
(A) Relative growth rate of *C. reinhardtii* with added co-culture exometabolomic extract without and with added thiamin (left) and with *M. aeruginosa* culture extract (right). Controls are with no addition or 15 μL methanol (+MeOH). (B) Scatter plot showing Orthogonal Projections to Latent Structures Discriminant Analysis (OPLS-DA) of peak areas for annotated metabolites detected via untargeted LC-MS in triplicate methanol extracts. (C) Volcano plot of metabolites in co-culture extracts relative to *C. reinhardtii.* Dots are calculated from triplicate measurements. Dotted line is *P* = 0.05. Thiamin antivitamins are shown as red dots. (D) Levels of thiamin antivitamins in *M. aeruginosa* single culture and co-culture extracts were measured by targeted LC-MS/MS using parallel reaction monitoring (PRM). For A and D data are means ± SEM, with n = 3. Asterisks indicate significance in Student’s t-test (\**P* < 0.05, \*\*\**P* < 0.001, \*\*\*\**P* < 0.0001, ns: not significant).

Volcano plots comparing the individual exometabolomes reveal that 891 and 86 compounds, common to both the co-culture to the single *C. reinhardtii* culture extracts, were present in significantly higher and lower amounts, respectively (Fig. 2*C*). A similar pattern was observed (839 metabolites in greater abundance versus 56 lower) when comparing the exometabolomes of singly cultured *M. aeruginosa* and *C. reinhardtii* (Fig. S3*A*). Many of the annotated chemicals are expected *M. aeruginosa* metabolites or metabolites observed in other microbial exometabolomes (27, 28).

Bacimethrin, a thiamin antivitamin precursor, was observed in all the original exometabolomes, but the levels were 730- and 640-fold higher, in the co-culture and single culture *M. aeruginosa* extracts respectively, than extracts of the *C. reinhardtii* medium (red dots in Fig. 2*C* and Fig. S3*A*). The chemical identity of bacimethrin was confirmed by comparing its MS/MS fragmentation pattern to that of a standard (Figs. S4 and S5*B*-*C*). Levels of bacimethrin measured during targeted LC-MS/MS were roughly 70% higher in co-culture extracts than in single culture extracts (Fig. 2*D*). Subsequent *C. reinhardtii* extracts, prepared in baked-clean glassware, contained no measurable bacimethrin (Fig. S5*A*).

Bacimethrin is known to be toxic to many bacteria and yeast (21–23). Its biosynthetic pathway was elucidated in *Clostridium botulinum* (21) where two of the three required genes are located in a gene cluster (21, 29) (thymidylate synthase and methyltransferase; Fig. 3*A* and *B*) whereas all three are collocated in some bacteria (30). In *M. aeruginosa*, two putative bacimethrin biosynthetic genes, thymidylate synthase and glycosyltransferase (also in a distinct gene cluster, Fig. 3*B*), and a third putative gene, a methyltransferase, were all up-regulated by 5- to 7-fold when grown in co-culture as compared to single culture (Fig. 3*C*). Gene upregulation correlated with higher levels of bacimethrin accumulation in the co-culture (Fig. 2*D*) supports an allelopathic role for bacimethrin production. We note the presence of the same or a similar gene cluster in other *Microcystis* spp. and that the thymidylate synthase in these clusters appears larger than in *C. botulinum* which may signify altered biochemical activity (Fig. 3*B*).

**Fig. 3.**
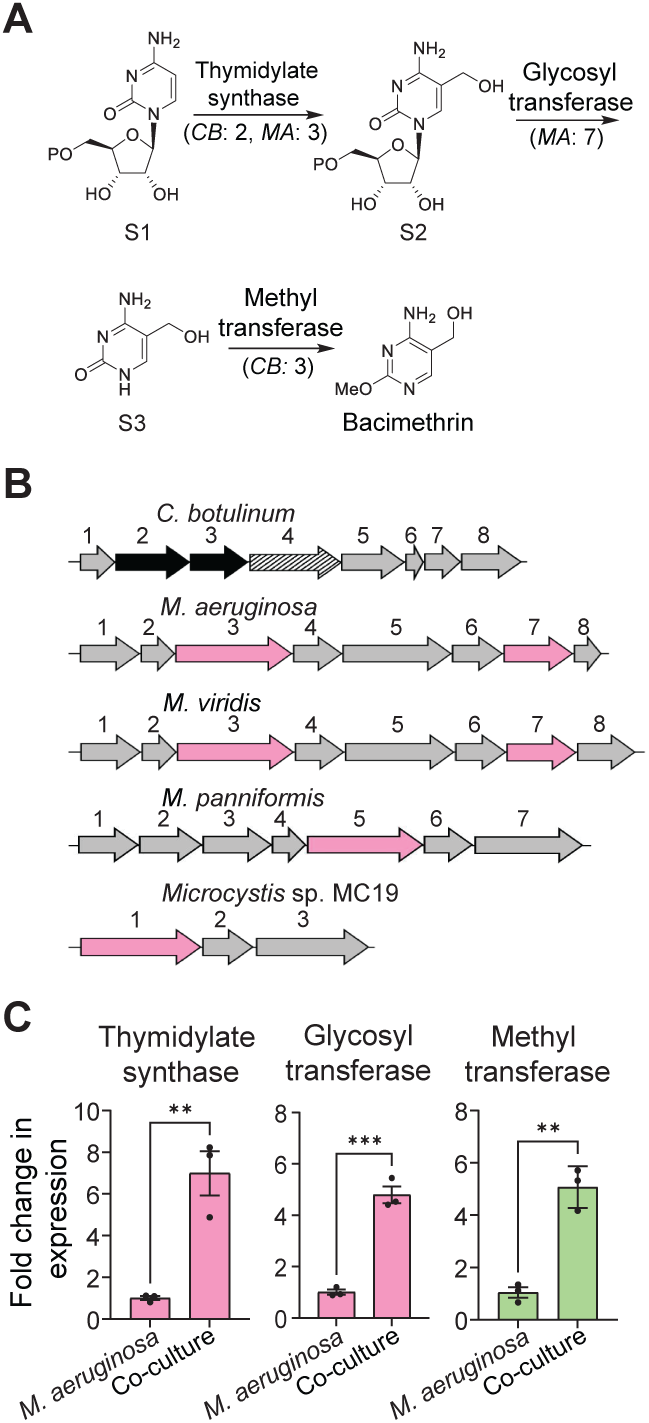
(A) Bacimethrin biosynthesis pathway adapted from (20) with gene number from cluster in B indicated in parentheses (*CB*: *Clostridium botulinum*; *MA*: *M. aeruginosa*). Substrate 1: cytidine 5′-monophosphate, 2: 5-hydroxymethylcytidine 5′-monophosphate, 3: 5-hydroxymethylcytosine. (B) Gene cluster for bacimethrin biosynthesis in *CB* strain A (ATCC 19397) and putative bacimethrin biosynthesis gene clusters in *MA* and three other *Microcystis* species (www.genome.jp/kegg/). In *CB*: 1- hypothetical protein (HP), 2- thymidylate synthase (TS), 3- methyltransferase (MT), 4- thiamin pyridinylase I, 5- phosphomethylpyrimidine kinase, 6- HP, 7- putative transporter, 8- ABC transporter. In MA and *Microcystis viridis*: 1- ABC transporter, 2- HP, 3- TS, 4- thymidylate kinase (TK), 5- mercuric reductase (MR), 6- phosphoribosyl-AMP cyclohydrolase/phosphoribosyl-ATP pyrophosphohydrolase, 7- glycosyl transferase, 8- HP. In *Microcystis panniformis*: 1- ABC transporter, 2- lipopolysaccharide transport system permease, 3- lipopolysaccharide transport system ATP-binding protein, 4-HP, 5- TS, 6- TK, 7- MR. In *Microcystis* spp. MC19: 1- TS, 2- pseudogene, 3- MR. (C) Relative expression of putative bacimethrin biosynthetic genes in *MA* determined by qPCR analysis. NCBI protein ID: thymidylate synthase (BAG04073), glycosyl transferase (BAG04077), and methyltransferase (WP_002744668). For C data are means ± SEM, with n = 3. Asterisks indicate significance in a Student’s t-test (\*\**P* < 0.01, \*\*\**P* < 0.001).

Methoxythiamin is formed when bacimethrin is entrained into the thiamin biosynthesis pathway via ThiD (31) (Fig. 4*A*), but it was not observed initially in our exometabolomes. However, using targeted LC-MS/MS and a synthesized standard [prepared following (32) with modifications, Fig. S6*A*], the presence of methoxythiamin was confirmed in the extracts derived from co-culture and *M. aeruginosa* single culture (Figs. S4 and S7). No difference between levels of methoxythiamin in the single culture and co-culture extracts was observed and extracted ion chromatogram (XIC) peak areas were roughly three orders of magnitude lower than those of bacimethrin (Fig. 2*D*); these results suggest that methoxythiamin is not the primary allelopathic chemical in our extracts although it is possible that methoxythiamin is transient in the culture medium perhaps because of chemical instability and/or rapid uptake by *C. reinhardtii*.

**Fig. 4.**
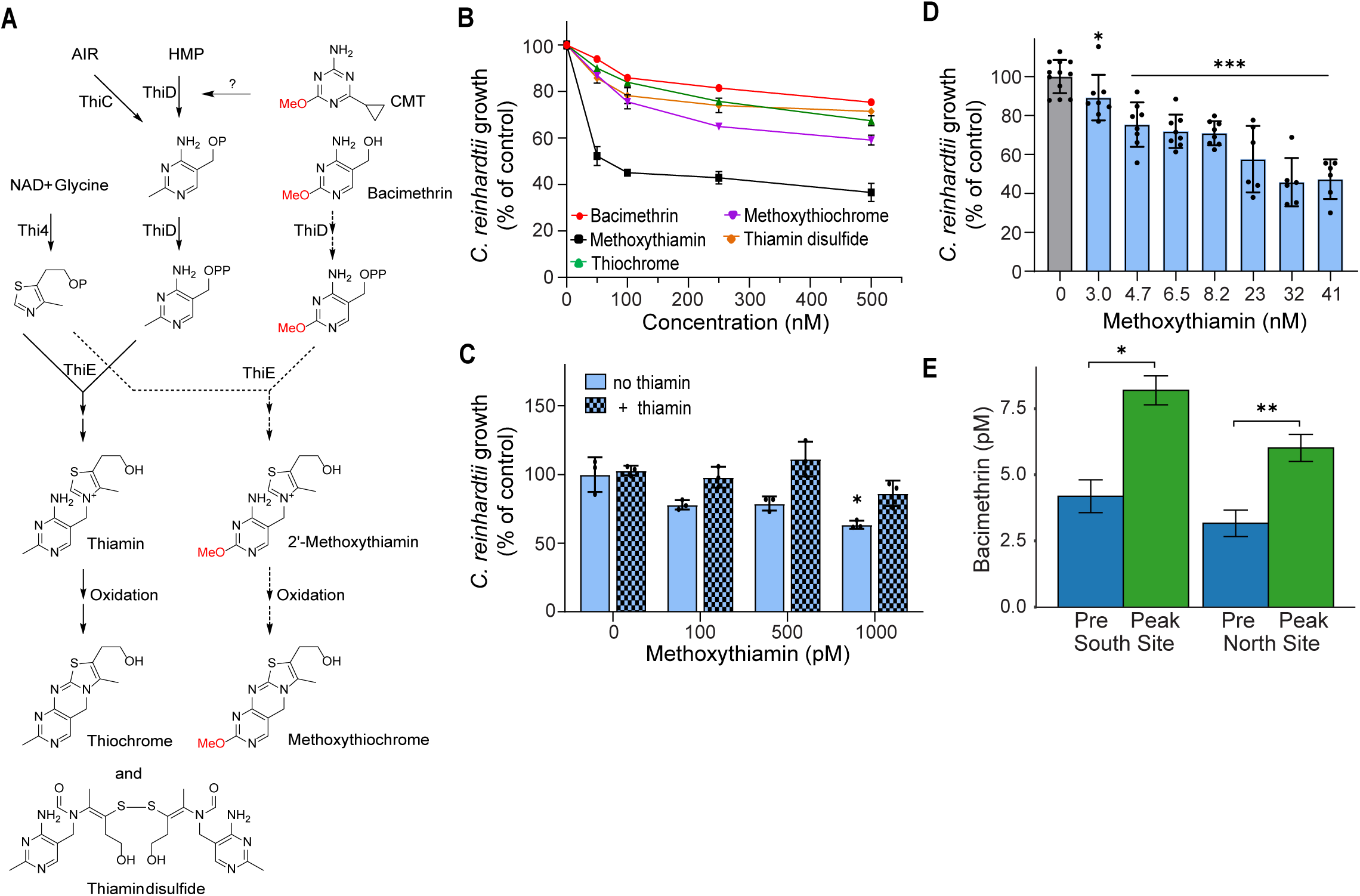
(A) Partial thiamin biosynthesis pathway for algae shown with known antivitamin integration steps (Ref 20), proposed inhibition by CMT, and structures of confirmed oxidation products. Enzyme and metabolite abbreviations: AIR, 5-aminoimidazole ribotide; HMP, 4-amino-5-hydroxymethyl-2-methylpyrimidine; ThiC, phosphomethylpyrimidine synthase; ThiD, HMP (phosphate) kinase; Thi4, thiazole synthase; ThiE, thiamin-phosphate pyrophosphorylase. (B) Average relative growth rates (n = 3) of *C. reinhardtii* with thiamin antivitamins added separately. Shown with thiamin additions in Fig. S9. (C) Average relative growth rates (n = 3) of *C. reinhardtii* in the presence of low methoxythiamin concentrations with no thiamin or with thiamin added at concentrations 10-fold lower than methoxythiamin (10, 50 and 100 pM respectively). Thiamin added to the control (no methoxythiamin) was 10 pM. (D) Average relative growth rates of wild-type *C. reinhardtii* exposed to methoxythiamin in microfluidic device. In (C) and (D) asterisks indicate statistically significant difference from replicated controls (0 nM) based on a Student’s t-test (in (C) \**P* < 0.008; in (D) \**P* < 0.025, \*\*\**P* < 0.002). (E) Dissolved bacimethrin concentrations in Upper Klamath Lake, Oregon. Bacimethrin was measured at the south and north ends of the lake (42.31046 N, 121.84369 W; and 42.46121 N 121.96013 W, respectively) in pre-bloom (blue bars, May 2023) and peak-bloom (green bars, August 2023) periods. Asterisks indicate statistically significant differences between sampling periods based on a paired t-test (\**P* = 0.011, \*\**P* = 0.0206). Bars are means (n = 3) ± StDEV.

Because thiamin can be transformed into the oxidation products thiochrome and thiamin disulfide via enzymatic or chemical reactions (33, 34), we hypothesized that methoxythiamin could likewise be oxidized to methoxythiochrome or methoxythiamin disulfide (Fig. 4*A*). Indeed, we found measurable levels of methoxythiochrome, as well as thiochrome and thiamin disulfide, in our exometabolomes at significantly higher concentrations in the co-culture and *M. aeruginosa* extracts compared to those of *C. reinhardtii* (Fig. 2*C* and Fig. S3*A*). Compound identity was confirmed using standards of thiochrome and thiamin disulfide (Fig. S4, MS/MS of thiochrome only in Fig. S8*A*) and we synthesized methoxythiochrome (Fig. S6*B*) to confirm its presence with targeted LC-MS/MS (Figs. S4 and S8*B*). Interestingly the methoxythiochrome XIC peak areas were 10-fold higher than methoxythiamin, but still 100-fold lower than those of bacimethrin, and like bacimethrin, levels were significantly higher in the co-culture extract (Fig. 2*D*). Thiochrome and thiamin disulfide peak areas were both roughly 10-fold lower than bacimethrin, but only thiochrome was higher in the co-culture extracts relative to *M. aeruginosa* (Fig. 2*D*). We found no evidence for methoxythiamin disulfide in any extracts.

### Thiamin Antivitamins and Oxidation Products Inhibit *C. reinhardtii* but not *M. aeruginosa*

We used batch cultures to confirm that the thiamin antivitamins were toxic to *C. reinhardtii* and also show that the various oxidation products were inhibitory. The chemicals were added individually to growth medium with and without supplemental thiamin. Methoxythiamin was the most toxic to *C. reinhardtii*; at the highest concentrations, the growth of *C. reinhardtii* was reduced by roughly 60% (Fig. 4*B*) and we saw significant inhibition as low as 1 nM (Fig. 4*C*). In comparison, methoxythiochrome was less inhibitory, only reducing growth by 40% at the highest concentration. Bacimethrin, thiochrome and thiamin disulfide were all similarly inhibitory, causing modest reductions in growth at the highest levels (∼15-25%, Fig. 4*B*). Adding excess thiamin (200 nM) reversed the inhibitory effect of all the thiamin-sized antivitamins except when the concentration of the antivitamin exceeded that of thiamin (Fig. S9*A*) and very low concentrations of thiamin (10-fold lower than antivitamin, 10 -100 pM) reversed inhibition at low methoxythiamin concentrations (Fig. 4*C*). In contrast, thiamin addition fully reversed the inhibitory effects of bacimethrin, even when bacimethrin was added at 60-fold higher concentration (Fig. S9*A*). It is likely that bacimethrin uptake by *C. reinhardtii* is facilitated by hydroxymethyl pyrimidine (HMP) transporters which may be down-regulated in the presence of thiamin, whereas the methoxythiamin and oxidation products are in competition with the added thiamin for uptake by thiamin transporters. Of these five chemicals, only the methoxythiamin was somewhat inhibitory to *M. aeruginosa* (∼25% growth reduction at 250 nM) and thiamin did not reverse inhibition (Fig. S9*B*). The primary source of the oxidized thiamin compounds is presumably *M. aeruginosa* since levels are similar in the co-culture and single culture extracts (Fig. 2*D*).

We also grew *C. reinhardtii* in the microfluidic device where we could maintain very low steady-state concentrations of the antivitamins. We observed statistically significant reductions in growth at methoxythiamin concentrations as low as 3 nM and reductions increased as concentrations increased, ultimately leveling out at about 50% of control (Fig. 4*C*), similar to batch culture experiments (Fig. 4*B*). We repeated batch and microfluidic device experiments with the *C. reinhardtii* thi8 mutant CC-25 (35), which lacks the ability to synthesize the pyrimidine moiety of thiamin. CC-25 cells were much more sensitive to both bacimethrin and methoxythiamin than wild-type. We observed a significant reduction in growth rate at 47 pM bacimethrin (compared to approximately100 nM in wildtype) and nearly complete inhibition as the concentration neared 1nM (Fig. 5*A*). In response to methoxythiamin, we saw a significant decrease in growth rate at concentrations as low as 15 pM, with complete cessation of growth occurring at 650 pM and higher (Fig 5*B*). As with the wildtype cells, very low concentrations of thiamin completely reversed inhibition in batch experiments (1-10 pM; Fig 5*C*).

**Fig. 5.**
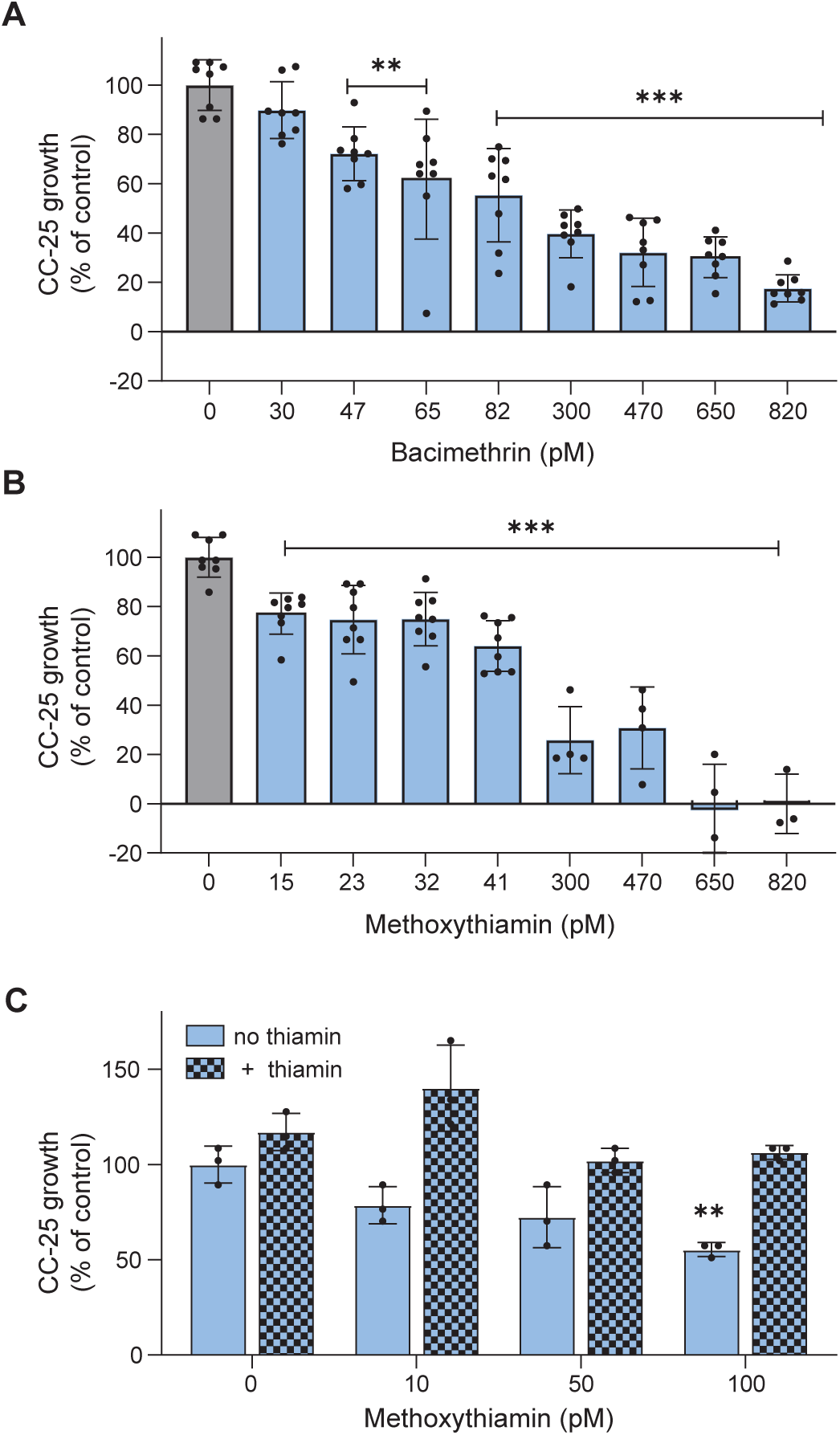
(A) Average relative growth rates of CC-25 exposed to bacimethrin in the microfluidic device. (B) Average relative growth rates of same exposed to methoxythiamin. (C) Average relative growth rates (n = 3) of CC-25 in the presence of low methoxythiamin concentrations in batch culture with no thiamin or with thiamin added at concentrations 10-fold lower than methoxythiamin (1, 5 and 10 pM respectively). Thiamin added to the control (no methoxythiamin) was 10 pM. In all three panels, asterisks indicate statistically significant difference from replicated controls (0 pM) based on a Student’s t-test (\*\**P* < 0.002, \*\*\**P* < 0.0001).

### Novel Antivitamin also Inhibits *C. reinhardtii*

Among the many chemicals elevated in the exometabolomes of *M. aeruginosa*, we confirmed the identity of one as 4-cyclopropyl-6-methoxy-1,3,5-triazin-2-amine (CMT) (Figs. S4 and S10 *A-C*). CMT is homologous to HMP and bacimethrin and was particularly abundant in both co-culture and *M. aeruginosa* exometabolomes (Fig S3*B*). Both the untargeted and targeted MS revealed significantly more CMT in the co-culture extracts than the *M. aeruginosa* extracts (Fig. S11*A*). CMT mirrored bacimethrin in its inhibition of *C. reinhardtii* and, likewise, thiamin addition eliminated its inhibitory effects (Fig. S11*B*). This suggests that, like bacimethrin, CMT is taken up via the HMP pathway and is perhaps inhibitory to ThiD, the enzyme that converts HMP to HMP-P (Fig. 4*A*) or other enzymes inhibited by bacimethrin (31). However, CMT is also structurally similar to the nonspecific triazine herbicide prometon which blocks electron transport in photosystem II (36). We could find no reference to biological production of CMT, though biosynthesis of cyclopropyl moieties is known (37), and we note that several other putative cyclopropyl-containing chemicals were annotated in the *M. aeruginosa* exometabolomes, including the known phytotoxin hypoglycin A, found in high levels in unripened lychee fruit (38). CMT was below detection in the *C. reinhardtii* extracts (Fig. S10*A*) and CMT did not inhibit *M. aeruginosa* (Fig. S11*B*).

### Bacimethrin is Present in Aquatic Environments Impacted by *Microcystis* Blooms

As an environmental validation of our laboratory investigation, we determined that dissolved bacimethrin was present in Upper Klamath Lake, Oregon, a hypereutrophic system that experiences annual cyanobacterial HABs (39–41). During the peak algal bloom period, observed bacimethrin concentrations were 8.2 ± 0.5 and 6.0 ± 0.5 pM at the south and north ends of the lake, respectively, roughly double those observed during the pre-bloom period (Fig. 4*E*). Bacimethrin has not previously been reported in the environment. These data provide evidence that at least one of the antivitamins described herein is present in the environment and further support our hypothesis that *M. aeruginosa* produces antivitamins to gain a competitive advantage.

## Discussion

We have documented here using co-culture cultivation and targeted MS that the HAB-forming freshwater cyanobacteria *M. aeruginosa* synthesizes the thiamin antivitamins bacimethrin and methoxythiamin. Toxic to a wide range of organisms (42), these chemicals are likely used by *M. aeruginosa* to compete with other photosynthetic algae. Production of these antivitamins by *M. aeruginosa* does not require co-culture but is substantially upregulated when a competing algal species is present. Furthermore, we found that thiamin and methoxythiamin oxidation products, as well as one novel compound with structural similarity to bacimethrin, all accumulated to higher levels in co-culture medium. Each inhibited the growth of wildtype *C. reinhardtii* when added individually with little or no effect on *M. aeruginosa*. Since the addition of thiamin largely eliminated the inhibitory effect of these compounds, it is likely that they all interfere with thiamin metabolism in *C. reinhardtii*. We also found that CC-25, a *C. reinhardtii* mutant unable to produce thiamin, is extremely sensitive to both bacimethrin and methoxythiamin. Also of note, our results may explain previous contradictory co-culture results: the green algae *Oocystis marsonii* was not inhibited by *M. aeruginosa* exudates when using WC medium (43) which contains thiamin whereas allelopathic interactions were noted between *Chlorella vulgaris* and *M. aeruginosa* when BG-11 was used and no thiamin addition was indicated (17).

We speculate that *M. aeruginosa* acquired the ability to synthesize bacimethrin from a bacterial source by horizontal gene transfer, a frequent occurrence in this species (44). It is noteworthy that both the *Microcystis* spp. and *C. botulinum* gene clusters contain an ABC transport gene (Fig. 3*B*) that is an ortholog of *YkoD*, the ATPase component of *YkoEDC*, a gene cluster believed to be involved in HMP transport in bacteria (45). The *M. aeruginosa* genome does not have copies of *YkoE* or *YkoC* but does contain *ThiXYZ* which has been linked to HMP uptake (45, 46). Given the ortholog’s sustained presence in the bacimethrin biosynthesis gene clusters, we speculate that it could be involved in the export of bacimethrin. The thiaminase I gene, present in the *C. botulinum* gene cluster (21), does not appear to have transferred to *Microcystis* spp. presumably because it would be of limited value, as an extracellular enzyme, to an organism living in a relatively dilute aquatic habitat.

We determined, for the first time, the presence of bacimethrin in a freshwater environment, Upper Klamath Lake, Oregon (Fig. 4*E*). This lake has an annually reoccurring *Microcystis* bloom that peaks in August, coincident with our sampling event (39–41). The concentration of bacimethrin in Upper Klamath Lake ranged from 3-8 pM and was roughly double at two sites during the algal bloom compared to pre-bloom, providing evidence that wild populations of *Microcystis* can produce bacimethrin. While the observed bacimethrin concentrations are quite low, they are within the published range of environmental concentrations (0.5-420 pM) and whole-cell uptake affinities (K_m_ 9.5 pM to 1.2 nM) of its non-toxic congener HMP (2, 19). Furthermore, the environmental bacimethrin concentrations close to those that caused inhibition in the thiamin-requiring mutant CC-25 (Fig. 5*A*). Additional research will be required to definitively connect environmental bacimethrin production to *Microcystis* ecophysiology and to determine the role that it and other antivitamins may play in structuring algal or perhaps even microbial communities.

While allelopathy has been argued against as a cause of HABs in marine ecosystems (18), it is more likely to play a role in freshwater ecosystems where cell densities are typically higher and turbulence is lower. Previous research on such interactions between *M. aeruginosa* and benign algae has primarily focused on the wide range of intracellular cyclopeptides produced by *M. aeruginosa* or lipophilic metabolites, though exact mechanisms of inhibition and their environmental significance are not fully understood (17, 47, 48). Antimetabolites from cyanobacteria with known inhibitory targets, such as the non-proteinogenic amino acid β-methylamino-L-alanine (BMAA)(49) or the shikimate pathway inhibitor 7-deoxy-sedoheptulose (50) are consequential in the context of cumulative trophic toxicity and drug discovery respectively, but they have not been shown to alter plankton composition. In contrast, thiamin antivitamins could effectively influence species composition because the concentrations of thiamin and its precursors can be extremely low in the water column (low to sub pM, 1, 2) and high affinity transporters are used in their acquisition (19, 45, 51).

Given thiamin’s scarcity and its critical function in universal biochemical pathways, researchers have long sought to understand the influence of thiamin on aquatic ecosystems and primary productivity (12, 13, 52, 53), but none previously addressed the role of antivitamins. Algae like *C. reinhardtii,* that are able synthesize thiamin (B_1_-prototrophs) but down-regulate biosynthesis in the presence of exogenous thiamin (54), likely take up these chemicals and may tolerate low levels. However, based on our experiments with the thiamin deficient mutant, organisms that are unable to synthesize thiamin (auxotrophs) will likely be very sensitive to antivitamins unless they have acquired resistance (55). While a relatively low percentage of eukaryotic algae are auxotrophic (3), percentages are higher amongst bloom forming species such as dinoflagellates, euglenoids and cryptomonads (5). These algal species may be the primary targets for the antivitamins. Reduced biomass of cryptomonads has been reported during a HAB bloom along with that of green algae and diatoms (56). It has also been reported that most naturally occurring bacterioplankton are auxotrophs for thiamin (4), and while *M. aeruginosa* is perhaps most often in direct competition with other phototrophs, its use of urea may lead to competition with heterotrophs (57).

These ecological findings, along with thiamin’s inherent instability in surface waters (including sensitivity to higher temperatures, 58), underscore the importance of gaining a deeper understand of this phenomena. Given the widespread prevalence and major impact of this toxin-producing organism on sensitive freshwater ecosystems worldwide, it is critical to understand its strategies for dominance. It is our hope that a better understanding of such mechanisms will lead to new strategies for control and abatement of HABs.

## Methods

### Algae Strains and Growth Conditions

Wild type *C. reinhardtii* (CC-125) and thiamin-requiring Thi-8 mutant (CC-25) were obtained from the Chlamydomonas Resource Center (University of Minnesota). *M. aeruginosa* PCC 7806 was purchased from the Pasteur Culture Collection (Paris, France). BG-11 (59) was used for experiments involving CC-125 and PCC 7806. BG-11 supplemented with 2% TAP and 1 nM thiamin was used for experiments involving CC-25. Further details of culture maintenance and experimental preparations are included in the SI Methods.

### Microfluidic Device

A microfluidic device with precise control of chemical environment was used to measure the growth rate both wildtype *C. reinhardtii* or CC-25 in an array of microhabitats (25). An agarose gel was molded with a silicon master and soaked in medium overnight. Cells are seeded onto the patterned gel which was then secured in a Plexiglas manifold atop a glass slide. Arrays of microhabitats are flanked by side channels through which source and sink medium are pumped with a syringe pump (Fig. 1*D*). Molecular diffusion through the agarose gel establishes a linear gradient of chemicals included in the source channel flow but absent from the sink channel (Fig. 1*D*). Details are included in SI Methods.

### Preparation of Exometabolome Extracts

*C. reinhardtii*, *M. aeruginosa*, and co-cultured cells were grown in BG-11 for 14 days to generate spent medium. A 0.2 μm syringe filter was used to remove cells and other debris. The filtrate was lyophilized in 50 mL polypropylene tubes and cellular metabolites were resuspended with 1 mL of methanol (MeOH).

### Chemical Syntheses

Methoxythiamin sulfate synthesis, based on a previous report (32) with some modifications, is shown in Fig. S6*A*. Methoxythiochrome synthesis is shown in Fig. S6*B*. Details are included in SI Methods.

### LC-MS Analyses

For untargeted analyses, exometabolome extracts were analyzed by ultra-high-performance liquid chromatography (U-HPLC, Thermo Vanquish) coupled to a quadrupole mass spectrometer QE-HF (Thermo) with an electrospray ionization (ESI) source in positive ion mode. For targeted analyses, the same HPLC was coupled to an Orbitrap QE-HF operated in positive ion mode using data dependent acquisition (DDA) and parallel reaction monitoring (PRM). The targeted CMT measurements were conducted on an Exion LC coupled with a X500B Q-TOF system. Details are included in SI Methods.

### Exometabolome Analysis

Orthogonal Projections to Latent Structures Discriminant Analysis (OPLS-DA) was used to analyze differences in the chemical composition of the exometabolomes obtained via untargeted analysis (n = 3 for each). Details of the OPLS-DA analysis are in SI Methods. Volcano plots were generated using GraphPad Prism 9. Compounds scoring below a threshold of *P* < 0.05 and exceeding a two-fold change either up or down (Log_2_ > 1 or Log_2_ < 1) were classified as significantly higher or lower respectively relative to the comparison exometabolome.

### RNA Extraction and qRT-PCR

RNA extraction from *M. aeruginosa* cells, cDNA synthesis, and qRT-PCR were done using standard protocols. Details are included in SI Methods.

### Environmental Bacimethrin Analysis

Samples for bacimethrin analysis were collected from two sites in Upper Klamath Lake (Oregon) using techniques previously described (1, 2). Sampling was conducted in May and August 2023 to capture the pre- and peak-bloom periods, respectively. Details are included in SI Methods.

## Data availability

All data generated for this paper is included in the figures, supplemental figures or in the supplemental data file (metabolomic data). If this paper is accepted for publication in mBio, we can deposit this data in the public repository MassIVE as necessary.

## Acknowledgments.

We thank Anurag Agrawal and Cliff Kraft for comments on the manuscript. We thank Kelly Shannon, Frederick Colwell, and Christie Nichols for assistance with environmental sample collection. This research was supported by the USDA National Institute of Food and Agriculture USDA Grant 67021-26598 (MW and BAA), US Fish and Wildlife Service Grant F22AC01810 (CPS), the California Department of Fish and Wildlife Q2196012 (CPS), Cornell University, College of Agriculture and Life Science (BAA) and Cornell University, College of Arts and Sciences (GWC).

## SI Appendix

### Methods

#### Algal Growth Conditions and Methods

All cultures (CC-125, wildtype *C. reinhardtii*, CC-25, thiamin-requiring Thi-8 mutant, and *M. aeruginosa* PCC 7806) were kept at 25 °C with 16 µmol m^-2^ s^-1^ continuous light unless otherwise stated. Batch cultures were placed on a shaker (130 rpm) in glass flasks (50 mL) or 96-well plates (300 µL). All experiments were initiated with exponential growth phase cells. Co-culture experiments were inoculated at cell densities of approximately 1 x 10^6^ cell mL^-1^ and 2 x 10^5^ cell mL^-1^ for *M. aeruginosa* and *C. reinhardtii* respectively. CC-25 cells were transferred twice into thiamin-free medium prior to experimentation to generate thiamin-limited cells for experiments. Cultures were received axenic, and we further confirmed that strains were axenic by visual examination and by adding 20 µL of each to Luria-Bertani (LB) bacterial medium; no bacterial growth was observed after 24 h (Fig. S12). Cell growth was monitored either by measuring cell density using a hemocytometer or chlorophyll *a* absorbance at 665 nm using a microplate reader (Power Wave XS, BioTek, USA). To minimize contamination in our exometabolomes, glass culture flasks were baked in a muffle furnace at 500 °C for 5 hours prior to use.

#### Batch Growth Inhibition Experiments

Wildtype *C. reinhardtii* and *M. aeruginosa* were grown into 96-well plates with additions of bacimethrin, methoxythiamin, thiochrome, methoxythiochrome, thiamin disulfide and CMT (0.10 - 500 nM) from stock solutions in MeOH. CC-25 was grown in 96-well plates with additions of methoxythiamin (10 -100 pM). Each was treatment was repeated with excess thiamin (200 nM) or less as specified in captions. Algal cell density was monitored daily with absorbance, and growth rate was calculated for each triplicate.

#### Microfluidic Device

As described in Kim et al. (1), algae were grown in microhabitats with defined chemical environments. Briefly, a silicon master with designed patterns was fabricated using SU-8 negative resist photolithography. A 3% agarose gel membrane was molded out of the silicon master and soaked in cell culture medium overnight before device assembly. For device assembly, 200 μL of *C. reinhardtii* cell culture at exponential growth phase (∼3 x 10^6^ cells mL^-1^) was first seeded onto the patterned agarose gel with four parallel sets of array microhabitats. The gel was then sandwiched between a Plexiglas manifold and a glass slide and screwed down to a stainless-steel frame to prevent leakage. Each set consists of a 4-by-4 array of 200 (L) × 200 (W) × 100 μm (H) microhabitats and two 400 (W) × 200 μm (H) side channels.

The flow through the side channels was controlled by a syringe pump (KDS230, KD Scientific, Holliston, MA) together with 10 mL syringes (BD, Franklin Lakes, NJ) and medical grade tubing (ID = 0.25 mm, PharMed BPT, Cole-Parmer, Vernon Hills, IL) which connected to the inlets of the microfluidic device using cut gel-loading tips. A flow rate of 0.7 μL min^-1^ was maintained throughout the experiment. The assembled device and the flow control unit were kept in an incubator at 25°C under 12 μmol m^-2^ s^-1^ illumination using fluorescent light bulbs for spent medium experiments (4200K, Lights of America; 3500K, SLI Lighting E-LITE) and under 100 μmol m^-2^ s^-1^ (Quantum Board, Horticultural Lighting Group) for the methoxythiamin exposure experiements. The entire setup was taken to the microscope for imaging once every day. Images were taken with an epi-fluorescence microscope (Olympus IX51, Center Valley, CA) equipped with a CCD camera (Cascade 512B, Photometrics, Tucson, AZ) and image acquisition software (IPLab imaging software, BD) was used to enumerate cell density. When exposing the cells to methoxythiamin (Fig. 4*C* and Fig. 5*C*), the source medium containing methoxythiamin (ranging from10 pM up to 50 nM) was replaced daily to reduce the impact of methoxythiamin degradation.

#### Chemical Synthesis of Methoxythiamin (MeOTh) and Methoxythiochrome (MeOTC)

Solvents for air sensitive reactions were purchased from Fisher, sparged with ultrahigh purity (UHP) grade nitrogen, and either passed through two columns containing reduced copper (Q-5) and alumina (THF) or passed through two columns of alumina (CH_2_Cl_2_) prior to use. All other chemicals and reagents were purchased from commercial sources (Sigma-Aldrich, Oakwood Chemical, Strem, TCI, Alfa Aesar, Acros, and Fisher) and used without further purification. All manipulations of air and water sensitive compounds were carried out under nitrogen by using standard Schlenk line technique. Flash column chromatography with silica gel (particle size 40– 64 μm, 230-400 mesh) was used to purify final products. ^1^H and ^13^C NMR spectra were recorded on Bruker AVANCE III HD (^1^H, 400 MHz) spectrometer with a BBF/1 H broadband observe probe or Bruker AVANCE III HD (^1^H, 500 MHz) spectrometer with a broadband Prodigy cryoprobe. All the NMR spectra were processed with MestReNova software. Chemical shifts (δ) for ^1^H NMR spectra were referenced to protons on the residual solvents (7.26 ppm for CDCl_3_, 3.31 ppm for CD_3_OD). Chemical shifts (δ) for ^13^C NMR spectra were referenced to the deuterated solvents themselves (77.16 ppm for CDCl_3_, 49.00 ppm for CD_3_OD). NMR spectroscopic data were reported as follows: chemical shift, multiplicity (s = singlet, d = doublet, t = triplet, br = broad), integration and coupling constants (Hz). High resolution mass spectrometry (HRMS) analyses were performed on a Thermo Scientific Exactive Orbitrap MS system equipped with an Ion Sense DART ion source.

MeOTh sulfate synthesis is shown in Fig. S6*A* and was based on previous report (2) with some modifications. In brief, for S4 synthesis, Et_3_N (0.04 ml, 0.28 mmol, 2.0 equiv) and CS_2_ (0.1 mL) was added to a solution of S1 (21 mg, 0.14 mmol, 1.0 equiv) in a mixture of THF (1 mL) and MeOH (0.5 ml) at 25 °C. After stirring for 10 min, a solution of S2(3, 4) (32 mg, 0.14 mmol, 1.0 equiv) in THF (0.5 ml) was added at 25 °C. The reaction mixture was further stirred for 30 min, then the solvent was removed under vacuum. The residual was redissolved in DCM (1 mL), and TFAA (0.03 mL, 0.20 mmol, 1.0 equiv) and Et_3_N (0.06 ml, 0.41 mmol, 3.0 equiv) was added at 25 °C. The reaction was further stirred for 1 h, then the solvent was removed under vacuum. The residual was redissolved in MeOH (1 mL), and 1 M aqueous KOH solution (1 mL) was added at 25 °C. The reaction was further stirred for 1 h, then diluted by water (10 mL) and further extracted with EtOAc (10 mL × 3). The organic phase was dried with Na_2_SO_4_, filtered, and concentrated under reduced pressure. The residual was purified by flash chromatography (DCM/MeOH = 40/1 to 20/1) to give S4 (20 mg, 47% yield) as a light-yellow solid. All the spectroscopic data matched with the literature report (2). For synthesis of MeOTh, a solution of aqueous H_2_O_2_ (1.0 M, 0.048 ml, 0.048 mmol, 3.0 equiv) was added to a suspension of S4 (5.0 mg, 0.016 mmol, 1.0 equiv) in aqueous HCl (0.27 M, 0.40 mL) at 0 °C. The reaction was stirred at 25 °C for 6 h and evaporated to dryness to give MeOTh (5.0 mg, 83% yield) as a white solid. All the spectroscopic data matched with the literature report (2).

For synthesis of MeOTC (Fig. S6*B*), MeI (0.17 mL, 2.7 mmol, 27 equiv) was added to a solution of S4 (31 mg, 0.10 mmol, 1.0 equiv) in MeOH (4 mL). After stirring at 60 °C for 4 h, the reaction mixture was cooled and all the volatiles were removed under reduced pressure. The residual was mixed with saturated aqueous NaHCO_3_ (5 mL) and extracted with DCM (10 mL × 3). The organic phase was dried with Na_2_SO_4_, filtered, and concentrated under reduced pressure. The remaining solid was washed with minimal amount of DCM to give MeOTC (7.5 mg, 27% yield) as a light-yellow solid. ^1^H and ^13^C NMR spectra of MeOTC dissolved in deuterated methanol (CD_3_OD) are shown in Fig. S13. MeOTC: ^1^H NMR (500 MHz, CD_3_OD) δ 7.93 (s, 1 H), 5.21 (s, 2 H), 3.90 (s, 3 H), 3.70 (t, *J* = 6.0 Hz, 2 H), 3.31 (s, 3 H), 2.78 (t, *J* = 5.9 Hz, 2 H), 2.23 (s, 3 H). ^13^C NMR (125 MHz, CD_3_OD) δ 171.78, 167.10, 164.09, 155.40, 133.88, 115.64, 104.39, 62.25, 55.08, 45.48, 30.36, 11.12. **HRMS** (DART-MS): m/z calculated for C_12_H_14_N_4_O_2_S^+^ [M+H^+^] 279.0911, found 279.0947.

#### LC MS Analyses of Methanol-Solubilized Exometabolomes

For untargeted analyses, lyophilized-algal medium methanol extracts were analyzed by ultra-high-performance liquid chromatography (Thermo Vanquish, UHPLC) coupled to a quadrupole mass spectrometer QE-HF (Thermo) operated with a electrospray ionization (ESI) source in both positive and negative ion mode. Positive ion mode was determined to be more effective for the compounds of interest. HPLC was performed with a Accucore Vanquish C18+ column (1.5 µm, 2.1 x 100 mm) heated to 30 °C using an injection volume of 2 µL. The metabolites were eluted via solute gradient at 200 µL min^-1^ using 0.1% formic acid in water (solvent A) and 0.1% formic acid in acetonitrile, ACN, (solvent B). The gradient was as follows: 0–2.0 min (0% B), 2.0–4 min (0–15% B), 4-14 min (15-32% B), 14-19 min (32-50% B), 19-19.1min (50-100% B), 19.1-21 min (100% B), 21-21.1 min (100-0% B), 21.1-23 min (0% B). Standards of bacimethrin (AdipoGen, San Diego, CA), methoxythiamin (synthesized as described), thiochrome (Sigma-Aldrich), methoxythiochrome (synthesized as described), thiamin disulfide (TCI America, Portland, OR), and 4-cyclopropyl-6-methoxy-1,3,5-triazin-2-amine (CMT, Alfa Chemistry, Protheragen Inc., Ronkonkoma, NY) were also run to confirm retention times observed in untargeted analysis. MS conditions were as follows: ESI voltage 3.5 kV, sheath, aux, sweep gas flow rates: 50, 10, 1 (arbitrary units), capillary temp. 275 °C, aux gas heater 375 °C, S-Lens RF level 55%. Software packages Compound Discovered 3.2 and Xcalibur 4.3, and Mass Frontier 8.0 were used for MS data analysis and prediction of in silico fragments.

For targeted analyses of the same extracts, the same HPLC and column was used at 55°C using an injection volume of 3 µL for samples and standards. Samples were diluted to 20% MeOH in 0.1% formic acid prior to injection to improve retention times; standards were prepared in same solution. The metabolites were eluted via solute gradient at 250 µL min^-1^ using 0.1% formic acid in water (solvent A) and 0.1% formic acid in ACN (solvent B). The gradient was as follows: 0–4.0 min (0.5-1.0% B), 4.0–8.5 min (1.0–20% B), 8.5-13.5 min (20-95% B), 13.5-15.5 min (95-99% B), 15.5-18 min (99-100% B), 18-19 min (100-0.5% B), 19-25 min (0.5% B). The HPLC was coupled to an Orbitrap QE-HF operated under positive ion data dependent acquisition (DDA) and parallel reaction monitoring (PRM) mode with targeted m/z for 5 standard precursors in the inclusion list. MS conditions were as follows: ESI voltage 3.8 kV, sheath, aux, and sweep gas flow rates: 20, 7, 1 (arbitrary unit), capillary temperature. 320 °C, aux gas heater 250 °C, S-Lens RF level 60. A sum of 5 to 7 product ions from each precursor with mass tolerance at 10 ppm was used to generate the XIC chromatograms in PRM data quantitative analysis by Xcalibur 4.3 software.

For the targeted CMT data included in the paper (Fig. S11*A*), the analysis was conducted on an Exion LC coupled with X500B Q-TOF system. A Luna C18 column from Phenomenex (3 µm, 2.0 x 100 mm) was used at a flow rate of 200 µL min^-1^, temp 30 °C. Solvents: A) 0.1% formic acid; B) 95% acetonitrile with 0.1% formic acid. A/B gradient: 0-5 min (5% B), 5-10 min (40% B), 10-13 min (90% B), 13-14 min (90% B), 14-15 min (5% B), 15-20 min (5% B). Injection volume: 2 µL for standards and 5 µL for samples, Loop volume 50 µL. MS analysis: Sciex X500B operated in ESI positive ion TOF mode. Calibration was carried out with negative calibrant using CDS system (ESI voltage: 5.5 kV; ion source gas1 and 2: 20 psi; curtain gas: 25 (arbitrary unit); CAD gas: 7 (arbitrary unit); source temperature: 350 °C; DP: 80V; accumulation time: 0.25 s; scan: an MS full scan from *m/z* 100 to *m/z* 1000 in profile mode followed by MRM HR scan acquired from 0 min to 20 min at CE 10V.

Sciex OS 2.0, was used for all data processing of targeted raw files generated in X500B, XIC chromatograms of five standards with mass tolerance at 0.02 Da from MRM HR runs are shown in Fig. S4.

#### Exometabolome Analysis

Orthogonal Projections to Latent Structures Discriminant Analysis (OPLS-DA) was used to analyze differences in the chemical composition of the exometabolomes using the *C. reinhardtii* data set as the control. The score plot for components t[1] and t[2] showed low variation among biological replicates (n = 3) and confirmed significant differences between the three separate algal exometabolomes (Fig. 2*B*). The two sample sets that include *M. aeruginosa* exudates scored negative values relative to t[1], but were distinct from each other along the t[2] axis, whereas the samples containing *C. reinhardtii* exudates were negative on the t[2] axis and distinct from each other along the t[1] axis. Axis values of t[1] and t[2] shown corrected by a factor of 1.00001. The R2X[1] was 0.753 and the R2X[2] was 0.188. The ellipse shown in the figure represents Hotelling’s T2 95% confidence range (Fig. 2*B*).

#### RNA Extraction and qRT-PCR

*M. aeruginosa* cells were pelleted by centrifugation from single cultures and co-cultures. RNA from triplicate pellets was extracted using the RNeasy Plant Mini Kit (Qiagen, Carlsbad, CA, USA). cDNA was synthesized from RNA using iScript^TM^ cDNA Synthesis Kit (Bio-Rad). Real-time RT-PCR was performed using gene specific primers (Table S1) and SYBR Green Supermix (Bio-Rad) in a CFX96 Real-Time PCR system (Bio-Rad). The *M. aeruginosa 16S rRNA* gene was used as internal control and relative changes in expression were analysed using the 2^-ΔΔCT^ method (5).

#### Environmental Bacimethrin Analysis

Samples for dissolved bacimethrin were collected from two sites in Upper Klamath Lake, Oregon, using techniques previously been described (6, 7); the southern site was offshore of Hanks Marsh (42.31046°N, 121.84369°W) and the northern site was at the mouth of the Williamson River (42.46121°N, 121.96013°W). Sampling was conducted in May and August 2023 to capture the pre- and peak-bloom periods, respectively. All sampling equipment was cleaned using 1 M hydrochloric acid and methanol as has previously been described (7). Samples were collected by boat from a depth of approximately 0.5 m using 1 L amber HDPE bottles (Nalgene). Samples were then prefiltered using a 100 micron mesh filter to remove metazoans, stored on ice, and transported back to shore for further processing. Gentle peristaltic filtration across a 0.2 micron Sterivex filter (PES membrane, Millipore, Burlington, MA, USA) was used to remove cells and suspended particles. The cell-free filtrate was acidified with 1 mL of 1 M hydrochloric acid and frozen at -20 °C until analysis. Sampling collection and processing in both May and August occurred mid-morning before local apparent noon.

Dissolved bacimethrin was extracted from lake water using solid phase extraction and its concentration was determined with liquid chromatography mass spectrometry (LC-MS) using a slight modification of previously described methods (6). Briefly, samples were thawed, pH adjusted to 6.5, and bacimethrin was extracted using C_18_ resin (Agilent Bondesil HF). Bacimethrin was eluted from the C_18_ resin using 12 mL of methanol, and further concentrated by nitrogen drying to a volume of 250 μL. A 1:1 chloroform liquid phase extraction was used to remove hydrophobic compounds from the sample matrix. Analysis was conducted using an Applied Biosystems 4000 Q-Trap triple quadrupole mass spectrometer with an ESI interface coupled to a Shimadzu LC-20AD liquid chromatograph. Chromatography and mass spectrometer conditions are described elsewhere (7). Similar to the analyses performed at Cornell, bacimethrin was observed to have column retention time of 2.89 minutes, a parent m/z of 156.1, and daughter product m/z of 138.0, 95.0, and 81.0. The declustering potential used was 20 V and the collision energy was 25 V. Samples were analyzed in triplicate and were randomized prior to analysis. An internal standard (^13^C-labeled thiamin) was used for quantification. Analysis was conducted at the Oregon State University Mass Spectrometry Center.

## Supplementary Materials

**Fig. S1.**
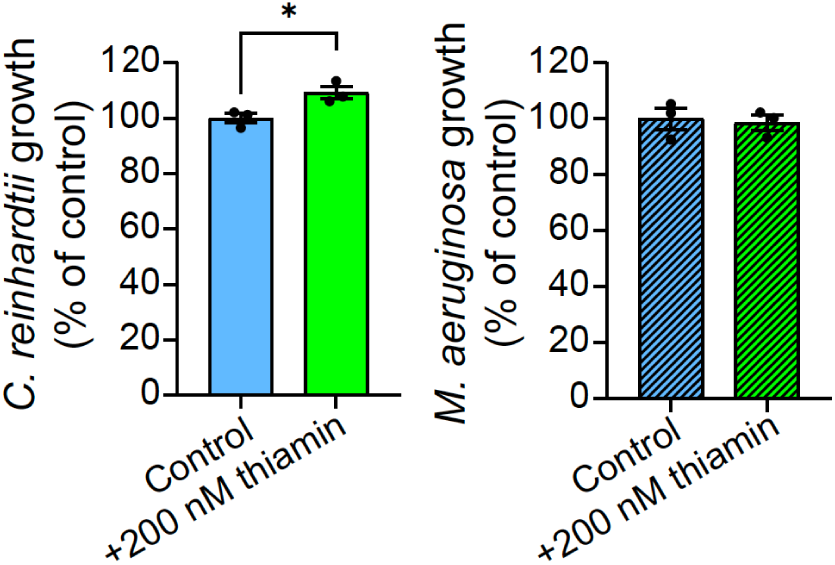
Thiamin addition is not inhibitory of *C. reinhardtii* and *M. aeruginosa* growth. Thiamin (200 nM) was added to the growth medium of both *C. reinhardtii* and *M. aeruginosa*. No adverse effect on growth rate was observed.

**Fig. S2.**
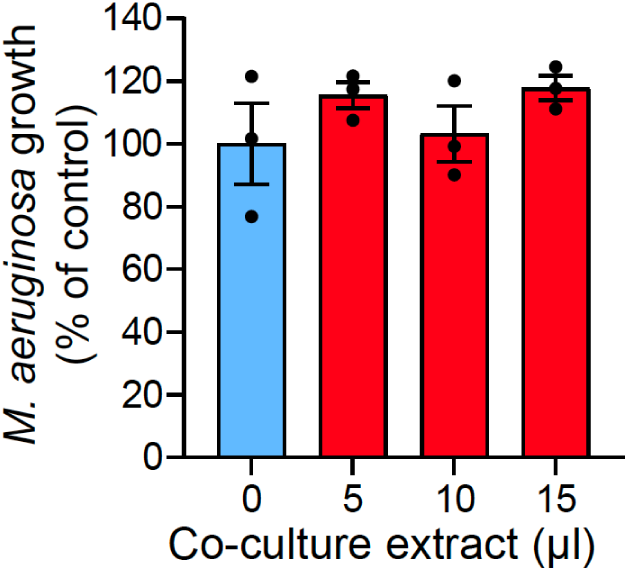
Methanol extract of co-culture spent medium does not inhibit *M. aeruginosa* growth. Relative growth rate of *M. aeruginosa* with methanol extract of co-culture medium added to growth medium.

**Fig. S3.**
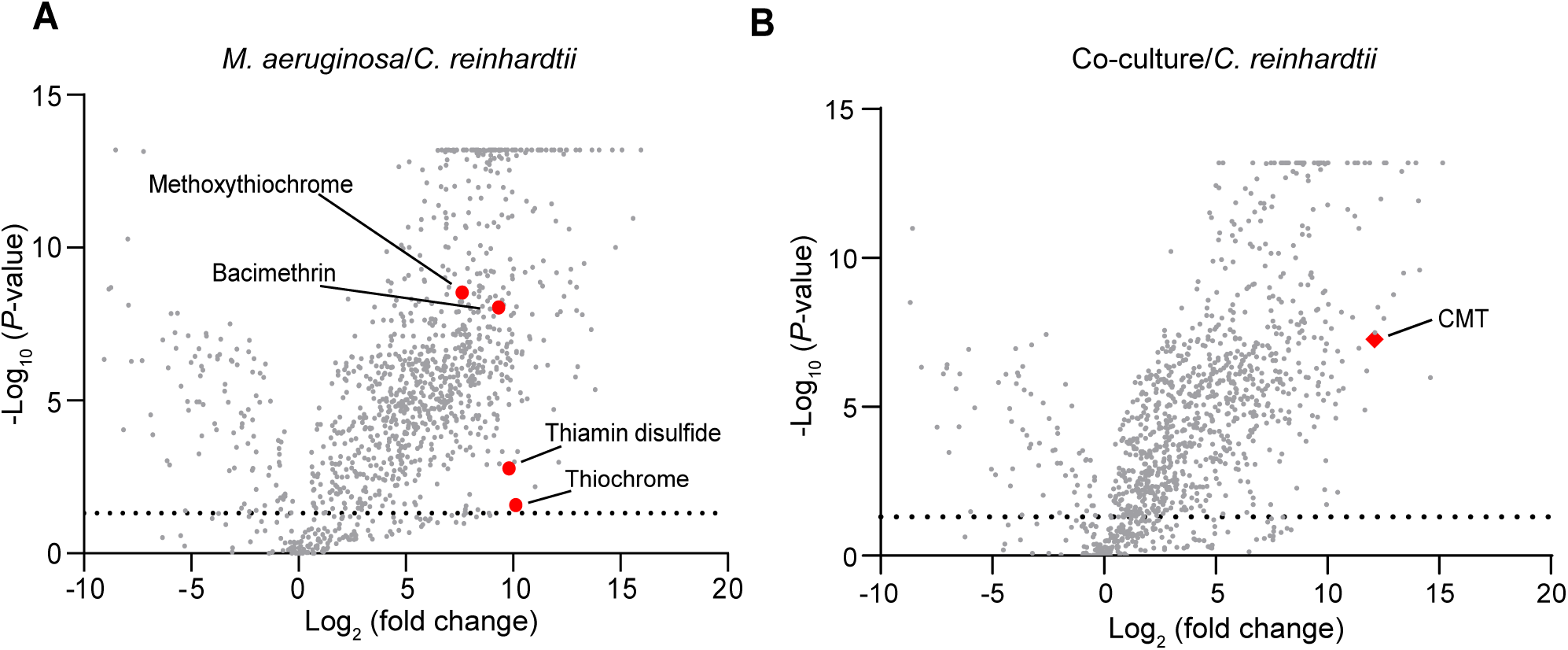
Thiamin antivitamin levels are many-fold higher in *M. aeruginosa* single culture and co-culture extracts compared to *C. reinhardtii* extracts. (A) Volcano plot of compounds found by LC-MS in *M. aeruginosa* extracts relative to *C. reinhardtii*. Thiamin antivitamins are shown as labeled red dots. (B) Volcano plot of compounds found by LC-MS in co-culture extracts relative to *C. reinhardtii.* CMT (4-cyclopropyl-6-methoxy-1,3,5-triazin-2-amine) is shown as labeled red diamond. In (A) and (B) dotted line: P = 0.05.

**Fig. S4.**
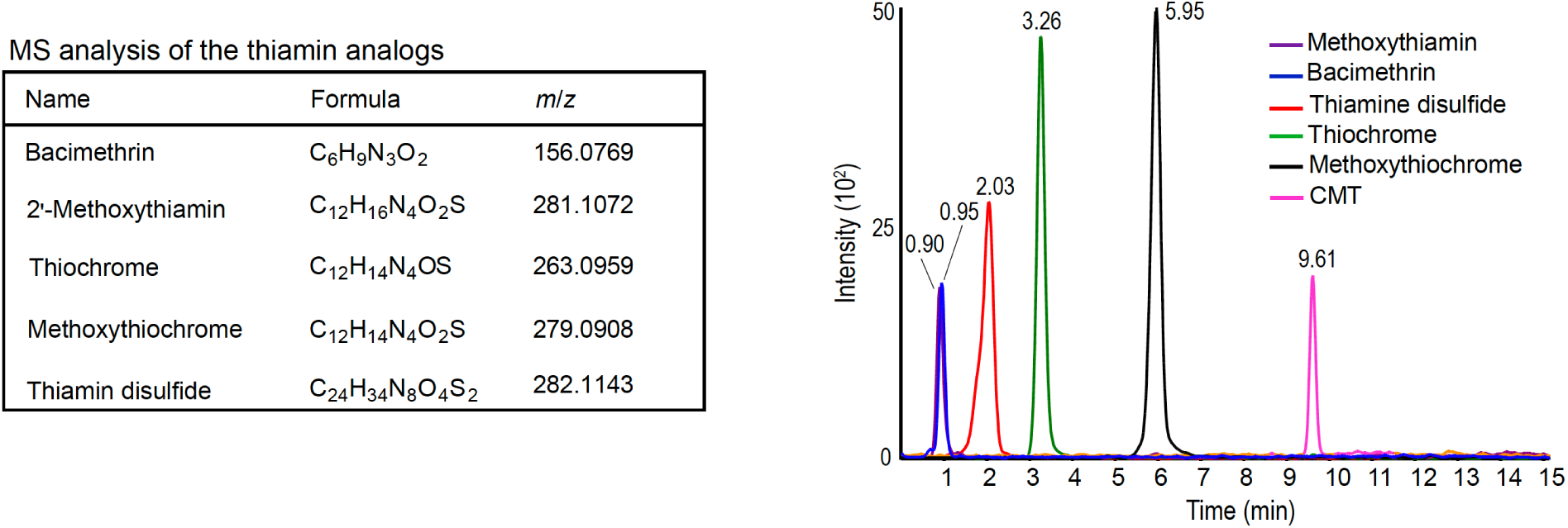
LC-MS/MS analysis of commercial and synthesized thiamin antivitamins. Names, formulas, and m/z of standards used. Composite chromatograms (XIC of PRM-HR) of the antivitamin standards labeled with retention time. Peaks shown were obtained by loading 40 pmol methoxythiamin, 1 pmol bacimethrin, 10 pmol thiamine sulfide, 5 pmol thiochrome, 15 pmol methoxythiochrome and 2 pmol CMT on a C18 column.

**Fig. S5.**
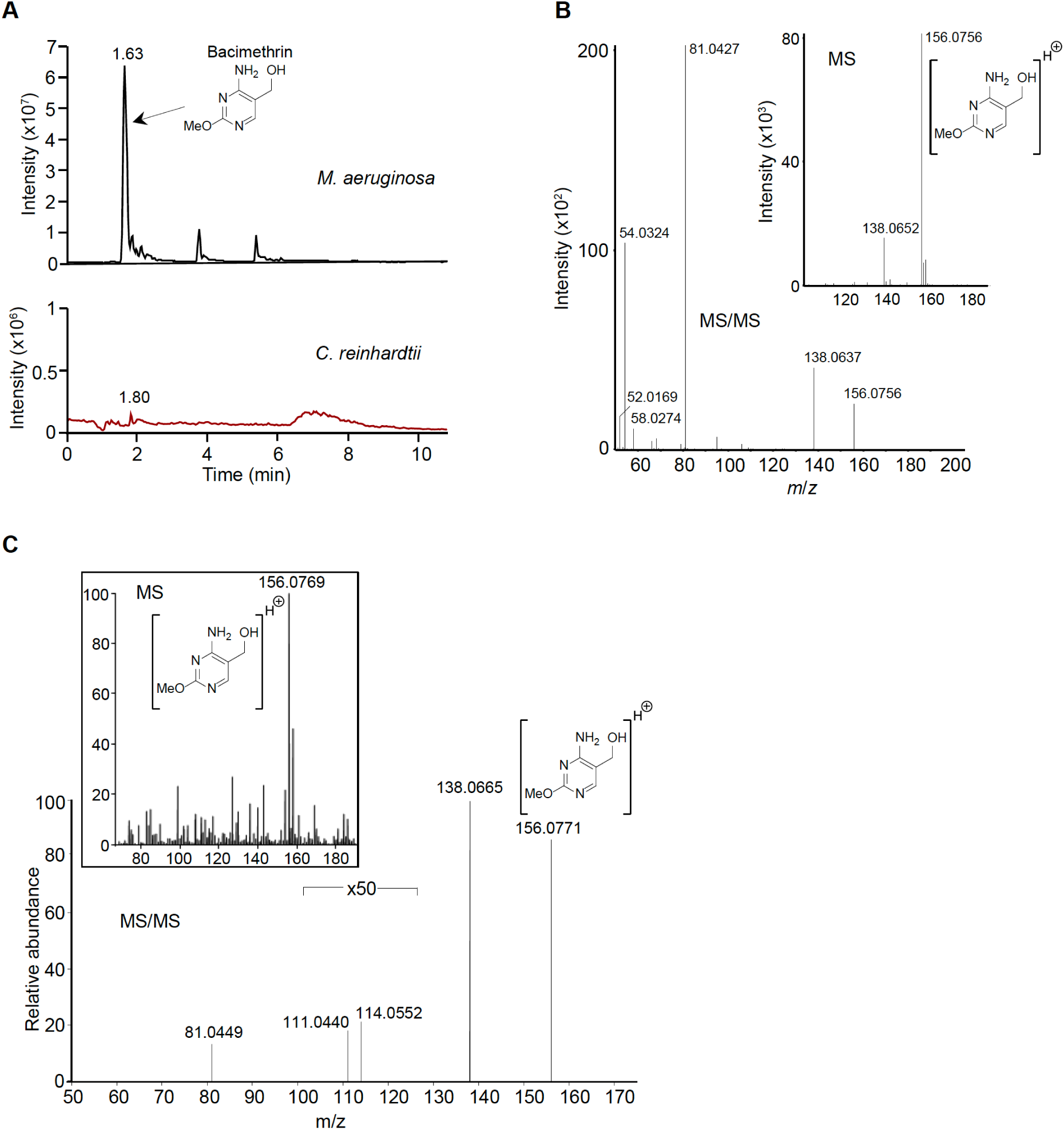
Presence of bacimethrin in *M. aeruginosa* extract is confirmed with LC-MS/MS. (**A**) Extracted Ion Chromatograms (XIC) of bacimethrin (m/z 156.0769) in *M. aeruginosa* and *C. reinhardtii* single culture extracts obtained by LC-MS/MS. (**B**) MS (inset) and MS/MS spectra of bacimethrin standard. **(C)** MS (inset) and MS/MS spectra of bacimethrin in *M. aeruginosa* single culture extract.

**Fig. S6.**
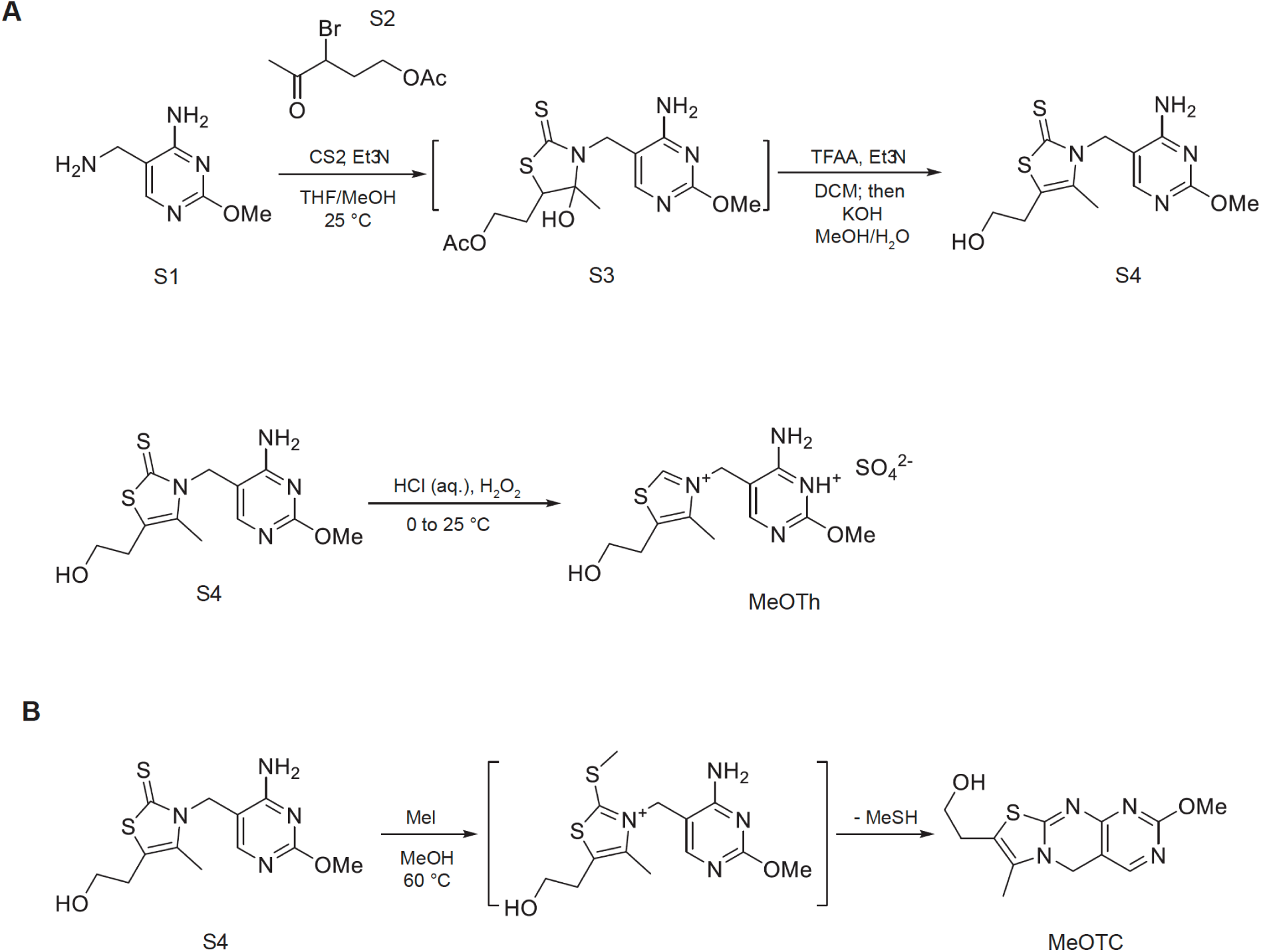
Chemical syntheses of 2′-methoxythiamin (A) and methoxythiochrome (B).

**Fig. S7.**
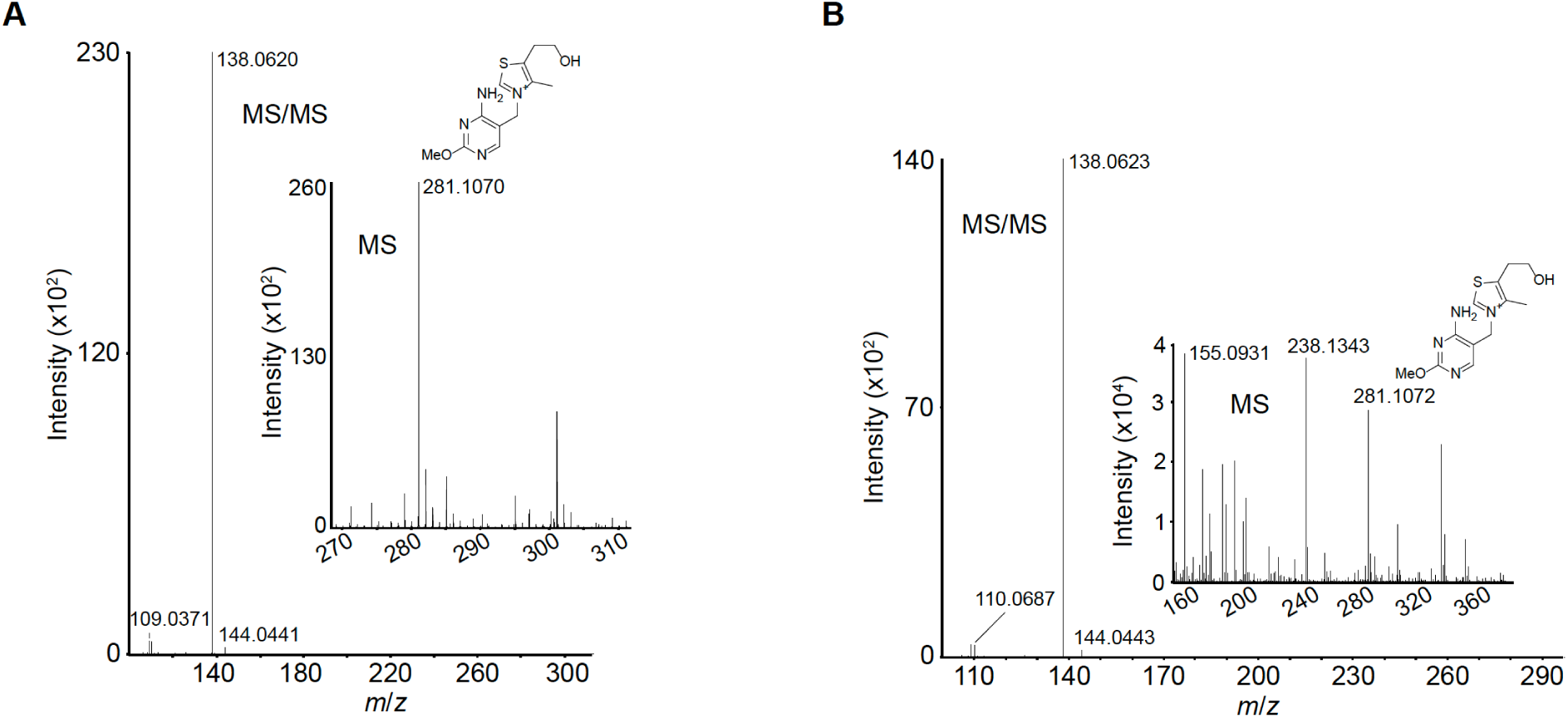
Presence of methoxythiamin in *M. aeruginosa* extract is confirmed with by LC-MS/MS. **(A)** MS (inset) and MS/MS spectra of methoxythiamin standard. **(B)** MS (inset) and MS/MS spectra of methoxythiamin in *M. aeruginosa* single culture extract. The MS/MS spectrum of m/z 281.106 for methoxythiamin was acquired using a relatively low collisional energy (CE = 15). The predominant fragment ion at m/z 138.065 was accompanied by several fragment ions with relatively low signal intensity.

**Fig. S8.**
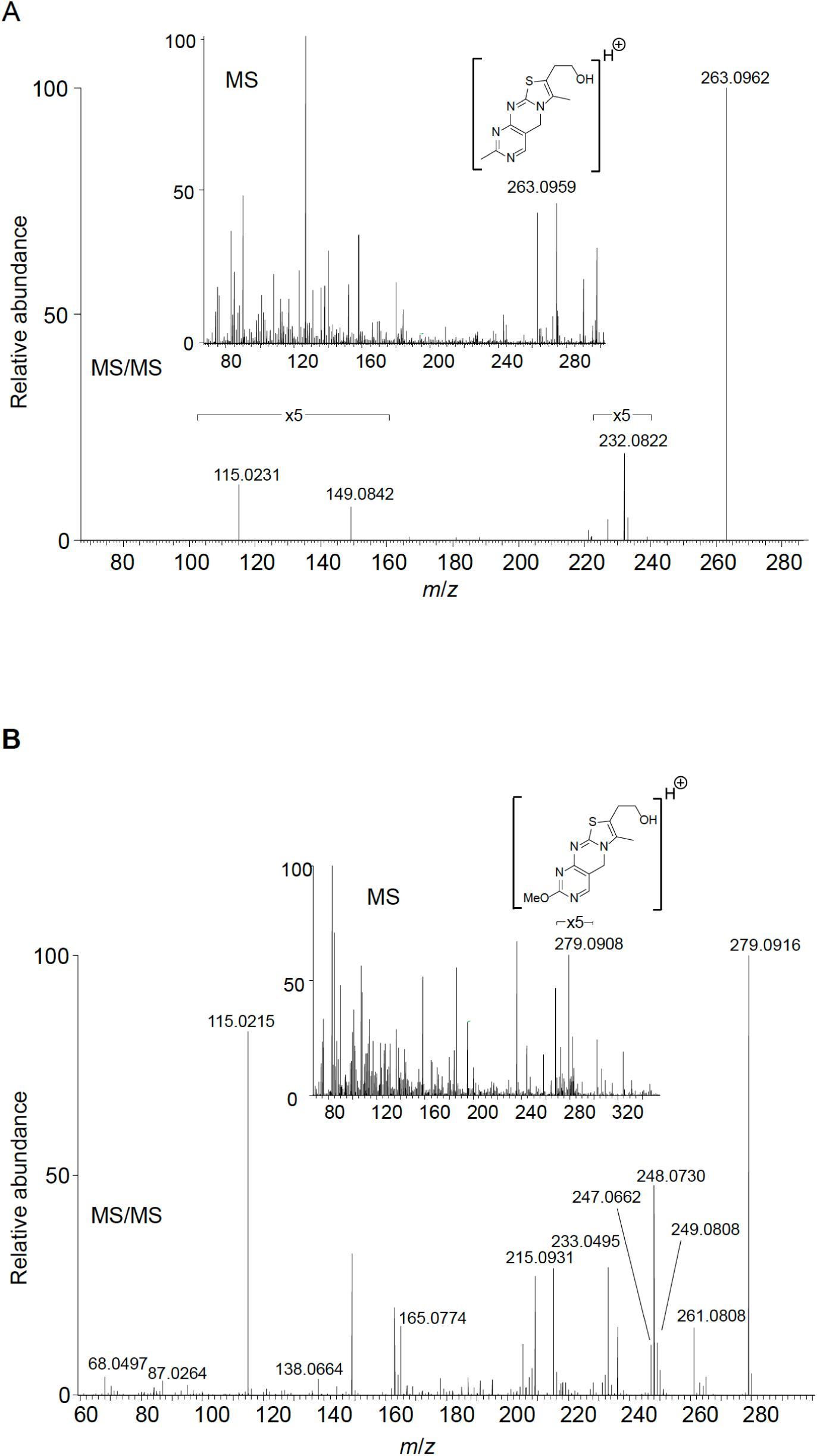
LC-MS/MS spectra for thiochrome (A)and methoxythiochrome (B). The chemical structure of the parent product is labeled on the MS spectra (inset) with the MS/MS spectrum.

**Fig. S9.**
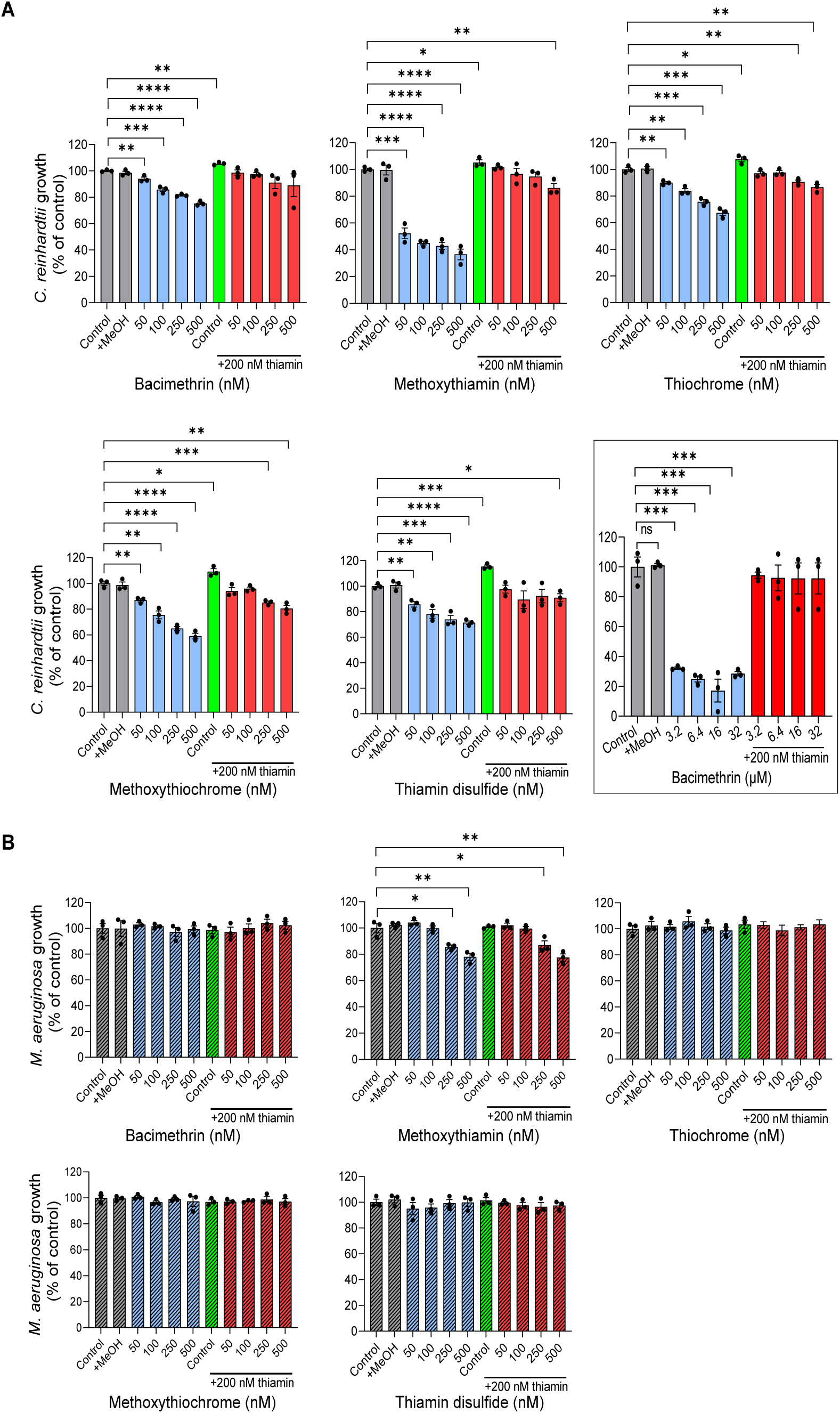
Thiamin antivitamins and their oxidation products cause thiamin deficiency in *C. reinhardtii* but not *M. aeruginosa*. **(A)** Relative growth rate of *C. reinhardtii* (relative to control with no added MeOH) with individual thiamin antivitamins added separately (blue bars) and added with 200 nM thiamin (green and red bars). Figure in box shows relative growth rate of *C. reinhardtii* with higher concentrations (µM) of bacimethrin. **(B)** Relative growth rate of *M. aeruginosa* (relative to control with no added MeOH) with individual thiamin antivitamins added separately (striated blue bars) and added with 200 nM thiamin (striated green and red bars). In (A) and (B) the volume of methanol (MeOH) added to the +MeOH control was equivalent to the highest volume of stock solution added. In (A) and (B) data are means ± SEM, with *n* = 3. Student’s *t* test was performed to show the significance of differences between data sets; \**P* < 0.05, \*\**P* < 0.01, \*\*\**P* < 0.001, \*\*\*\**P* < 0.0001.

**Fig. S10.**
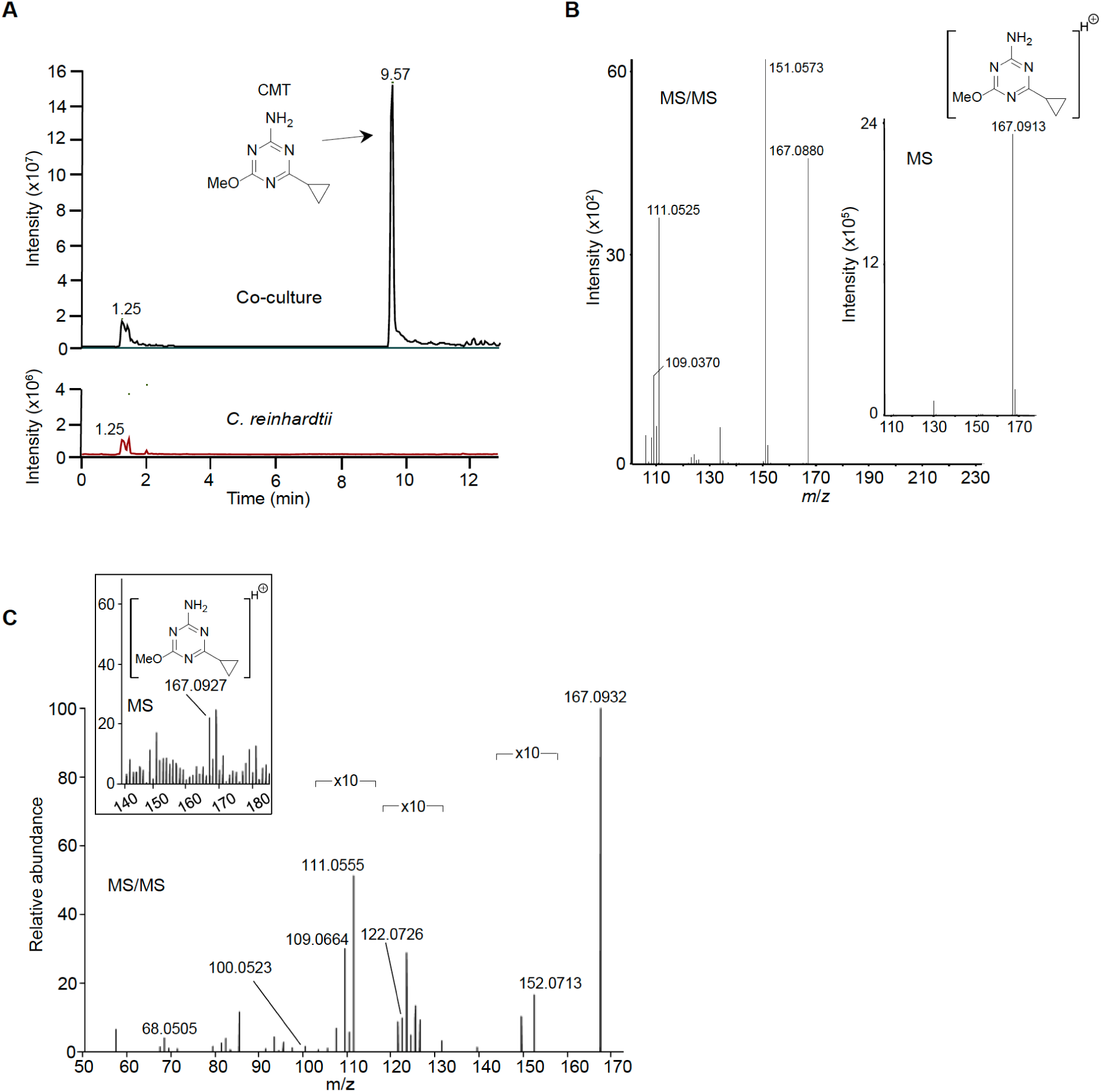
Presence of CMT in *M. aeruginosa* extract is confirmed with LC-MS/MS. **(A)** Extracted Ion Chromatograms (XIC) of CMT (m/z 167.0927) in co-culture and *C. reinhardtii* single culture extracts obtained by LC-MS/MS. (**B**) MS (inset) and MS/MS of CMT standard **(C)** MS (inset) and MS/MS of CMT in *M. aeruginosa* single culture extract.

**Fig. S11.**
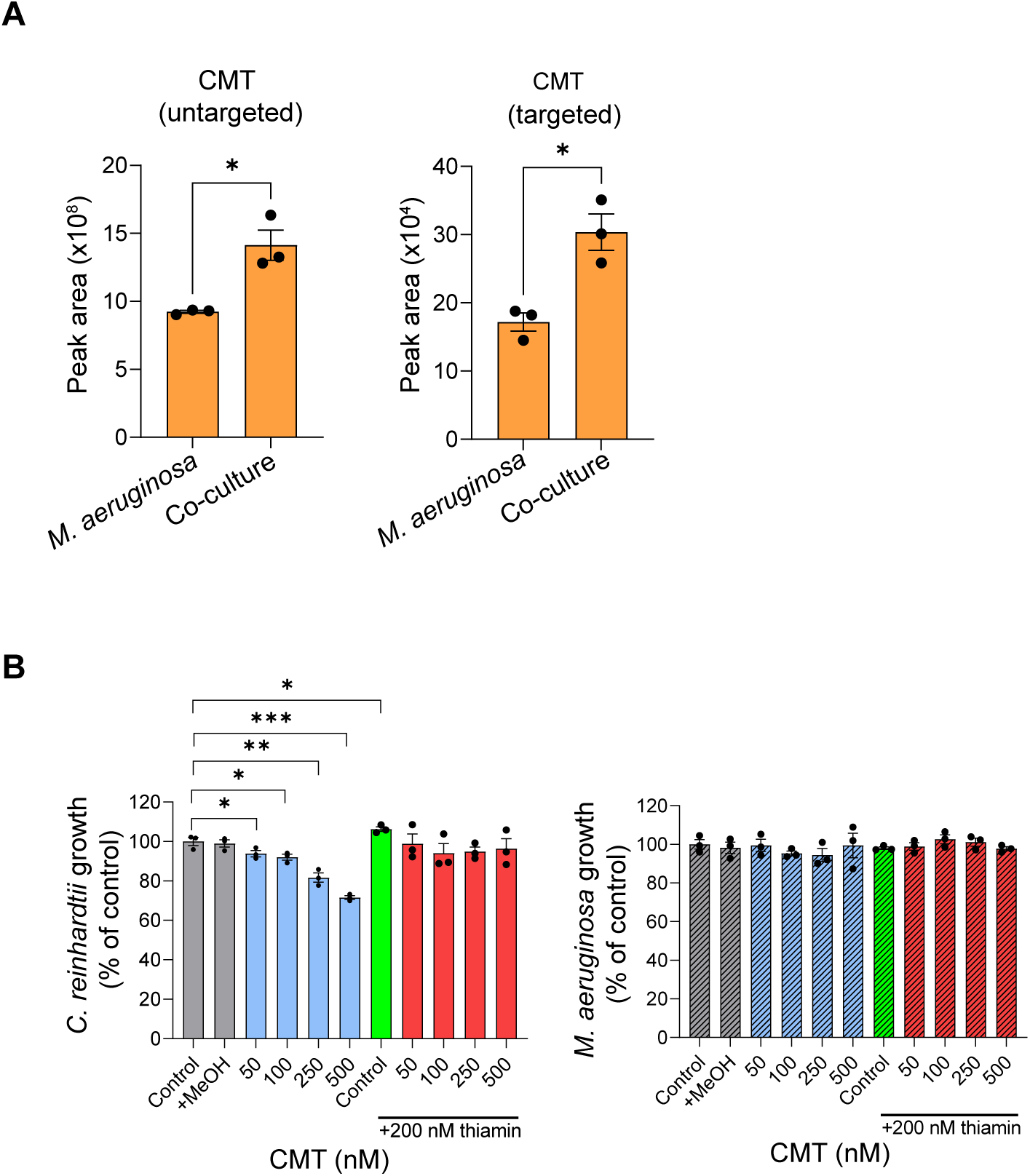
HMP-like analog CMT is elevated in co-culture exometabolome and causes thiamin deficiency in *C. reinhardtii* but not *M. aeruginosa*. (A) Levels of CMT in *M. aeruginosa* single culture and co-culture extracts obtained by nontargeted (left) and MRM LC-MS/MS (right). (B) Relative growth rate of *C. reinhardtii* (blue bars) and *M. aeruginosa* (blue striated bars) with added CMT. Green and red bars show relative growth rates with 200 nM thiamin added. Rates are relative to control with no added MeOH. The volume of MeOH added to the +MeOH control was equivalent to the highest volume of stock solution added. In (A) and (B) data are means ± SEM, with n = 3. Student’s t test was carried out to show the significance of differences between data sets; *P < 0.02, **P < 0.01, ***P < 0.001, ****P < 0.0001.

**Fig. S12.**
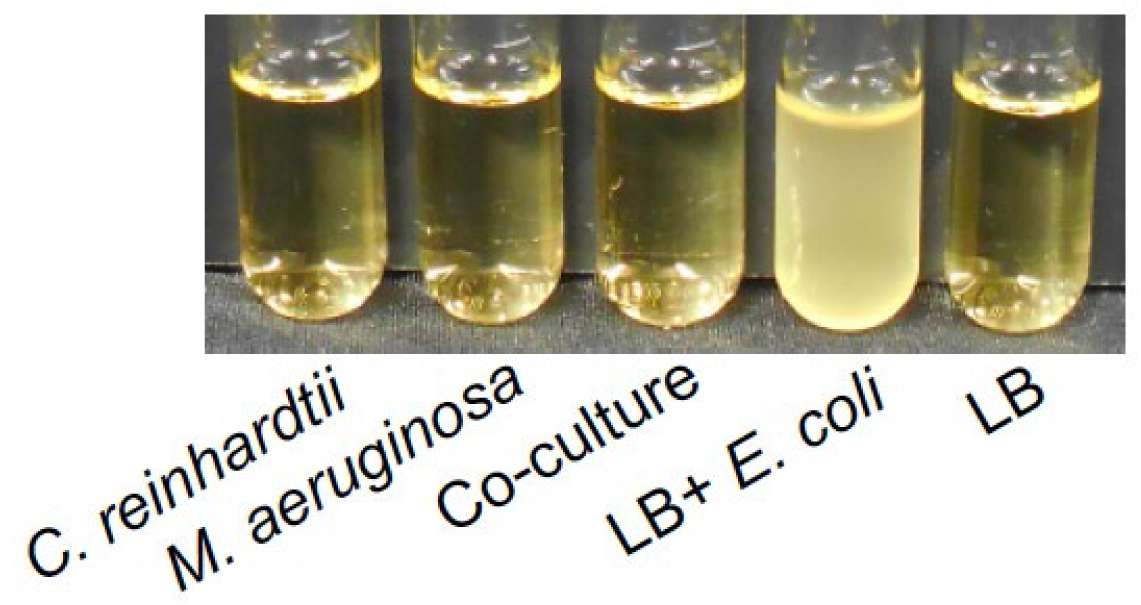
Visual confirmation that algal cultures are not contaminated with bacteria. A small volume (20 µl) of cell-free medium from individual cultures and co-culture was added to bacterial growth medium (LB medium) in glass culture tubes. Tubes were incubated overnight in a 37 ^◦^C shaker. LB medium showed no signs of bacterial growth unless inoculated with *E. coli*.

**Fig. S13.**
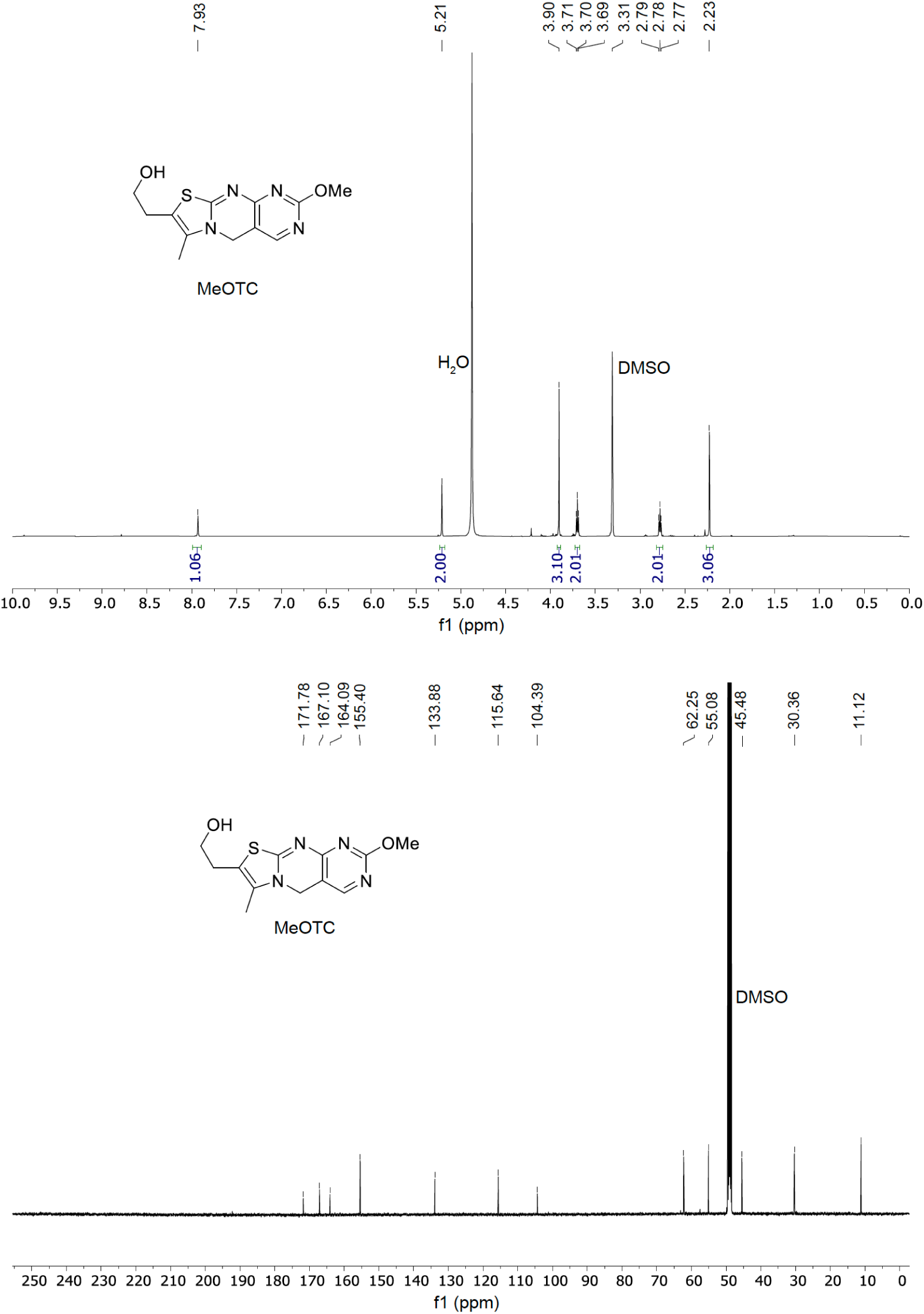
NMR spectra of methoxythiochrome (MeOTC). ^1^H NMR of MeOTC was measured at 500 MHz in deuterated methanol (CD_3_OD) (top) and ^13^C NMR of MeOTC was measured at 125 MHz also in CD_3_OD (bottom).

**Table S1:**
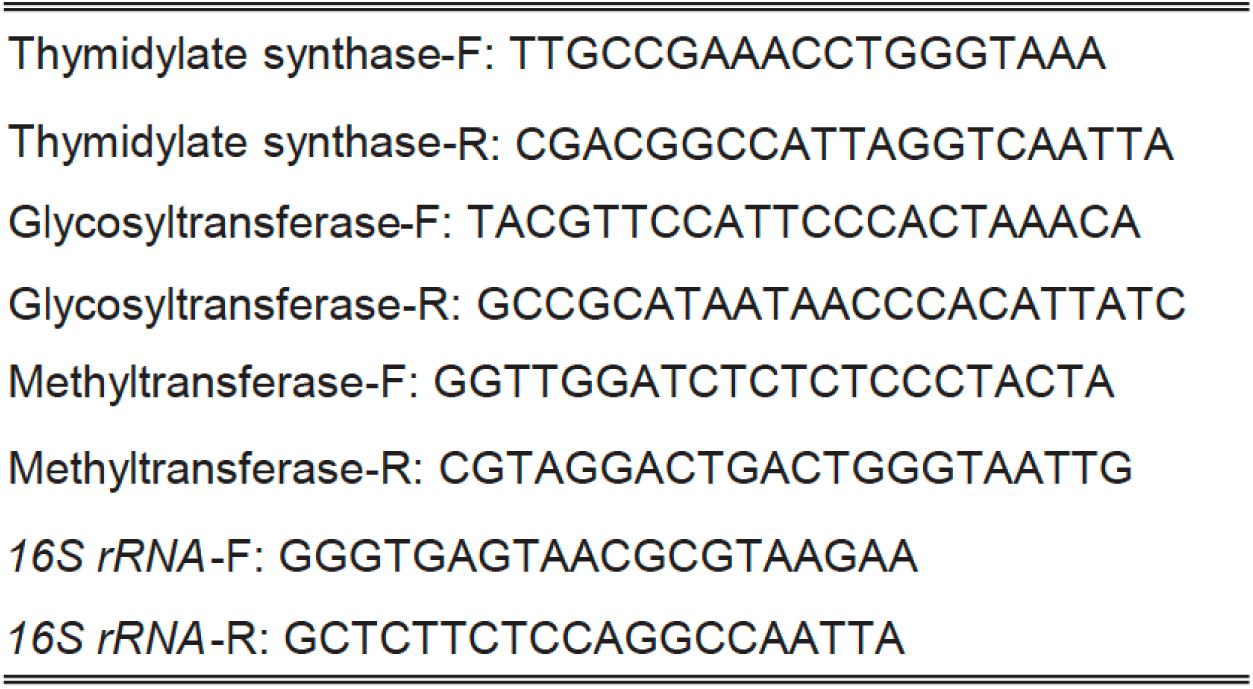
Primers used in this study for qRT-PCR of the bacimethrin biosynthetic genes in *M. aeruginosa*.

